# Semantic redundancy-aware implicit neural compression for multidimensional biomedical image data

**DOI:** 10.1101/2023.08.22.554284

**Authors:** Yifan Ma, Chengqiang Yi, Yao Zhou, Zhaofei Wang, Yuxuan Zhao, Lanxin Zhu, Jie Wang, Shimeng Gao, Jianchao Liu, Xinyue Yuan, Zhaoqiang Wang, Binbing Liu, Peng Fei

## Abstract

With the rapid development in advanced imaging techniques, massive image data have been acquired for various biomedical applications, posing significant challenges to their efficient storage, transmission, and sharing. Classical model-or learning-based compression algorithms are optimized for specific dimensional data and neglect the semantic redundancy in multidimensional biomedical data, resulting limited compression performance. Here, we propose a Semantic redundancy based Implicit Neural Compression guided with Saliency map (SINCS) approach which achieves high-performance compression of various types of multi-dimensional biomedical images. Based on the first-proved semantic redundancy of biomedical data in the implicit neural function domain, we accomplished saliency-guided implicit neural compression, thereby notably improving the compression efficiency for large-scale image data in arbitrary dimensions. We have demonstrated that SINCS surpasses the alternative compression approaches in terms of image quality, compression ratio, and structure fidelity. Moreover, with using weight transfer and residual entropy coding strategies, SINCS improves compression speed while maintaining high-quality compression. It yields near-lossless compression with over 2000-fold compression ratio on 2D, 2D-T, 3D, 4D biomedical images of diverse targets ranging from single virus to entire human organs, and ensures reliable downstream tasks, such as object segmentation and quantitative analyses, to be conducted at high efficiency.

## Introduction

Advanced imaging techniques in conjunction with efficient image processing approaches makes big impact on modern life science. Many biomedical applications currently require a vast amount of experimental data to be generated for various image-based analysis. For instance, studying the cytotoxic mechanisms of CAR-T cells through long-term live-cell imaging of cell morphological changes can produce several terabytes (TB) to tens of terabytes image data using high-resolution and high-throughput fluorescence microscopy systems ^1, 2^. For another instance, volumetric imaging of a mesoscale mouse brain at single-cell resolution to create a whole-brain neuron connectivity map will yield tens of terabytes image data^3^. Such a vast amount of image data places significant burdens on data storage, computation and sharing. For storage at limited space, these massive raw data have to be saved partially, with increased risk of data loss. Besides, due to the limited transmission bandwidth, researchers have to transfer and share data in an inefficient manner. Meanwhile, in contrast to centralized cloud storage and exchange technologies, it is crucial to achieve effective and data-specific storage compression directly at the user terminal. To this end, storage optimization on the data generation source^4^ and more importantly, on the downstream compression side should be studied.

The essence of compression lies in the removal of redundancy brought by the internal correlation among signals. Traditional compression methods based on analytical or statistical model explicitly remove spatial and temporal redundancy through transformations and coding, such as domain transformations ^5^ and entropy coding ^6, 7^, to substantially compress the data. Besides the spatial and temporal redundancies, there are also plenty of semantical similarities— for example, visually-similar cells in a microscopic image or a video containing the dynamic changes of the same target, which are widely existed in diverse biomedical image data. Experiments reveal that within these visually similar images, there is also a form of redundancy which is different from classic temporal and spatial redundancies, and termed as semantic redundancy ^8, 9^. It’s difficult for traditional model-based methods to capture these relatively abstract semantic correlations, thus leaving ample space for further enhancing the compression efficiency. Meanwhile, classic compression methods, such as JPEG ^10^, H.264 ^11^, H.265 ^12^ are designed for natural images/videos and thus perform poorly in compression of high dynamic range biomedical images. Moreover, frequency-domain-based compression methods may suffer from spectral truncation or blocking artifacts ^13^, which affects the accuracy of downstream analysis tasks. Therefore, for biomedical images characterized by high dimensions (time, 3D space, spectrum), high dynamic range, and high structural similarities, conventional compression techniques often show limited fidelity insufficient for subsequent quantitative analyses.

Unlike model-based approaches, deep learning-based data compression techniques, such as Autoencoders ^14^, VAE ^15^, GAN ^16^ *etc*. have recently emerged to interrogate the semantic correlation among signals. These approaches compress input data into a low-dimensional space representation which aims to eliminate semantic redundancy among data information by learning deep feature representations, and then reconstruct the original data using a decoder. Nevertheless, capturing essential data features, reducing dimensionality for optimal latent representation, and handling the burden of training with massive data for a single decoder remain highly challenging. Furthermore, when dealing with complex biomedical data, these deep learning-based supervised methods exhibit limited generalization capabilities and significant performance degradation due to generalization errors. The latest development of implicit neural representation (INR) ^17^ utilizes a neural network to parameterize a continuous function based on the data dimensions, enabling advanced representations of 3D scenes ^18^, images ^19^, and videos ^20^. Unlike traditional CNN-based approaches that utilize discrete, grid-like representations of image data, the grid-free feature of INR representation naturally fits with the continuity of target visual information. This facilitates the generation of continuous representations that enable seamless and arbitrary interpolation of visual data. In addition, with the universal approximation theorem of neural networks ^21, 22^, INR implemented with Multi-Layer Perceptron (MLP) can fit any complex function with a sufficient number of parameters, resulting in high-fidelity compression representation. INR currently has been demonstrated to be applicable for compressing natural image scenes ^23^ as well as various multi-dimensional biomedical data for biomedical research and clinical diagnosis, including 2D images, 3D volumes ^24^, and 4D data. By controlling the network parameters across different samples, stable compression rates and compression quality can be achieved. Moreover, INR exhibits stable performance for various data compression without the requirement of modifying the network structure specifically, which is can’t be achieved by other supervised learning-based approaches due to the generalization errors.

However, existing INR-based compression approaches require massive training time ^25-27^ due to the processing of a large number of input coordinates when representing compression data by network optimization. As the dimensions increase, the network parameters of INR also grow exponentially, resulting in significantly increased computation ^28^.Also, these INR approaches can’t effectively leverage the correlations between data, such as the inter-frame correlation in dynamic biomedical data and highly similar local features. Furthermore, INR fits the entire image region without specific optimization for particular signals. These challenges prevent the INR-based compression from surpassing the alternative techniques and also spurred the development of our new INR approach to overcome the limitations.

In this study, based on our first-proved semantic redundancy of biomedical data in implicit neural function domain, we propose the Semantic redundancy based Implicit Neural Compression guided with Saliency map (SINCS) approach, which explores the weight clustering effect in the implicit neural function domain and substantially accelerates the compression time of the algorithm through weight transfer. We also introduce saliency-guided compression mechanism to adaptively capture the specific structure information, thus facilitating high-fidelity compression of multi-modal biomedical images, and design residual-based entropy coding to further compress the optimized INR weights. Taken together, SINCS efficiently achieves high-fidelity compression with a high ratio up to 2000 folds for diverse multidimensional images. We have demonstrated SINCS’s superior compression performance on several large-scale biomedical image sets (2D, 2D video, 3D and 3D video) obtained from different imaging techniques (optical microscope, electron microscope, CT), proving its strong potentials for advancing a broad range of biomedical applications.

## Results

### High semantic correlation in implicit neural function domain of biomedical imaging data

The essence of compression is to eliminate the redundancy caused by the internal correlation among signals. Most of the existed compression methods only explore and eliminate temporal and spatial redundancy among data, ignoring semantic redundancy, which is common in multidimensional data, such as 2D picture, 2D video, 3D video, *etc*. The features in multidimensional biomedical images, such as organelles in the time-lapse video of a live cell, neurons in different regions of a large brain tissue image, are also correlated with semantic redundancy at both spatial and temporal dimensions. Here, we used zebrafish embryo heart as target to classify the biomedical image data into three modes for analyzing the sematic redundancy at different dimensions. In Mode 1, we analyzed the semantic redundancy along lateral dimensions using 2D plane image of the embryo heart. We employed the “patch sliding” to obtain the small patches from whole zebrafish embryo heart. Then we selected adjacent patches for compression using INR. By computing the Wilson Coefficient (Method section) to measure the distribution discrepancy of all the network parameters, we validated distributional correlation between the patches. In the domain of Implicit Neural Function (INF), the central tendency of the distribution is much more similar, whereas in the spatial domain, it presents a multi-modal distribution, as comparatively shown in Fig. 1a. The lower Average Wilson Coefficient (AWC) values calculated in INF domain, as compared with those in spatial domain, indicate much stronger sematic correlations found in INF domain. With discrepancy distribution and AWC metrics, we also analyzed the semantic redundancy along axial dimension (Mode 2) and temporal dimension (Mode 3) using the 3D image stack and 4D image video of the embryo heart, respectively. The results (2^nd^ and 3^rd^ rows) are consistent with those from lateral dimension (1^st^ row), validating that biomedical images with arbitrary dimensions are all suited for being represented and compressed in the INF domain of INR. In addition to the correlation measurement of all network parameters, we calculated correlation of the different layers, and found that the hidden layers showed relatively high correlation (Supplementary Fig. 1).

**Figure 1.**
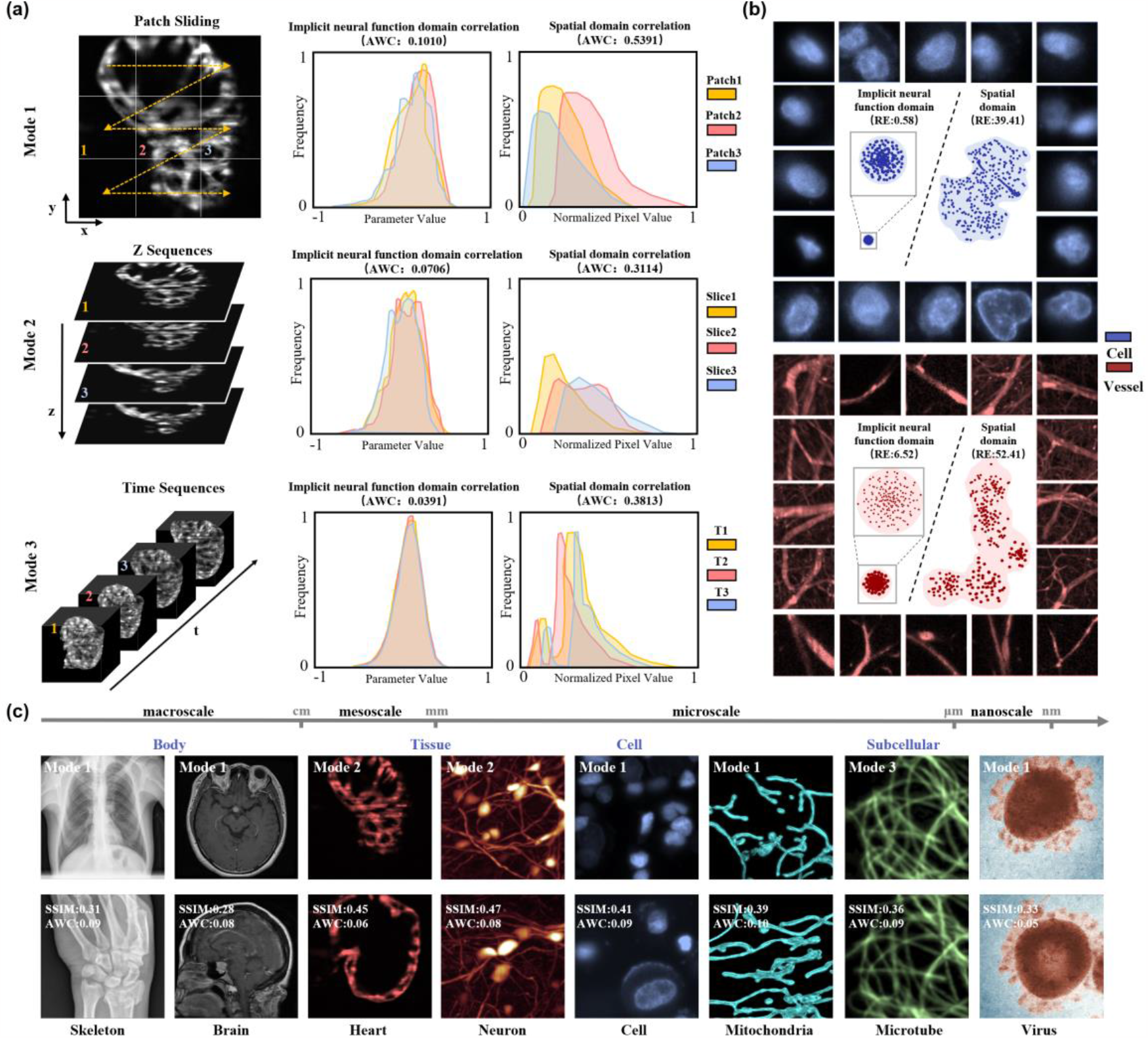
Validation of semantic redundancy in implicit neural function domain correlation. **(a)** Comparison of semantic correlations in three image modes. 2D, 3D and 4D (3D+T) images of the same zebrafish embryo heart were used as target, to evaluate the lateral (mode 1, top), axial (mode 2, middle), and temporal (mode 3, bottom) semantic correlations, respectively. In each mode, the images’ semantic correlations in the implicit neural function domain and spatial domain are compared through calculating their parameter histograms. The Average Wilson Coefficient (AWC) values are used as correlation metric with lower value indicating higher correlation. **(b)** Clusterings for structurally-similar data in implicit neural function and spatial domain. Blood vessels and cell nuclei are chosen to represent line-like and point-like signals, respectively. The Rayleigh Entropy (RE) values are calculated to quantify the clusterings, with lower value indicating more compact clustering of the images. **(c)** Comparison of image correlations in implicit neural function domain (AWC metric), and in spatial domain (SSIM metric). Multi-scale samples captured by different imaging techniques are compared to validate the universally-high correlations in implicit neural function domain. The human skeleton, human brain images are obtained from CT (Multi-Slice Spiral CT, Medium slices with 2.5mm thickness) and MRI (T1-weighted MRI), respectively; the 3D images of zebrafish embryo heart and mouse brain neurons are obtained by light-sheet microscope (4×/0.13 NA illumination and 20×/0.5 NA detection for heart, 4×/0.28 NA illumination and 10×/0.3 NA detection for mouse brain neurons); the 2D subcellular images of cell nuclei and mitochondrial are captured by light-sheet microscope (20×/0.45 NA detection objective for cell nuclei and 60×/1.1 NA detection objective for mitochondrial), and the 4D subcellular images of dynamics microtubes are obtained from single objective light-sheet microscope (100×/1.5 NA for illumination and detection); the virus images are obtained from electron microscopy (an LEO (Zeiss, Oberkochen, Germany) with a Morada (Olympus) camera).

We further explored this internal correlation in INF domain on typical point-like (cell nuclei) and line-like signals (blood vessels), which belong to basic components in most of the biomedical data. Through “block partitioning” of large-scale data, we generated three thousand patches containing local features from different spatial positions or time points. Then we applied T-distributed Stochastic Neighbor Embedding (t-SNE)^29^ dimensionality reduction (Method section) to compare the clustering patterns of these images in both the spatial domain and the INF domain (Fig. 1b). In sharp contrast to the muti-model distributions in the spatial domain (right part), very unimodal distributions were found in the INF domain (left part). The Rayleigh Entropy (RE) was calculated to quantitatively evaluate their clustering characteristics (Method section). The significantly lower RE values indicated a much more compact clustering of massive samples in INF domain, also suggesting that different samples of the same type could be rapidly compressed using Meta-learning through weight transfer in the INF domain. Furthermore, such high semantic correlation represented by low AWC and RE metrics in INF domain also proved wide existence in arbitrary-dimensional images (2D, 2D-T, 3D, 4D) of diverse biomedical targets (virus, organelles, cells, animal tissues, Human organs) with different size (nano-, micro-, meso-, macro-scale) and from different imaging techniques (electron microscope, light microscope, CT, MRI), as shown in Fig. 1c. More comprehensive quantifications are provided in Supplementary Note 1 and Supplementary Fig. 2 and 3 to cross-validate this universal INF domain correlation inside and between biomedical images. In the following step, we designed saliency map-informed and meta-learning-enabled SINCS to fully utilize such implicit semantic correlations to realize high-fidelity, high-ratio compression with improved speed.

**Fig. 2.**
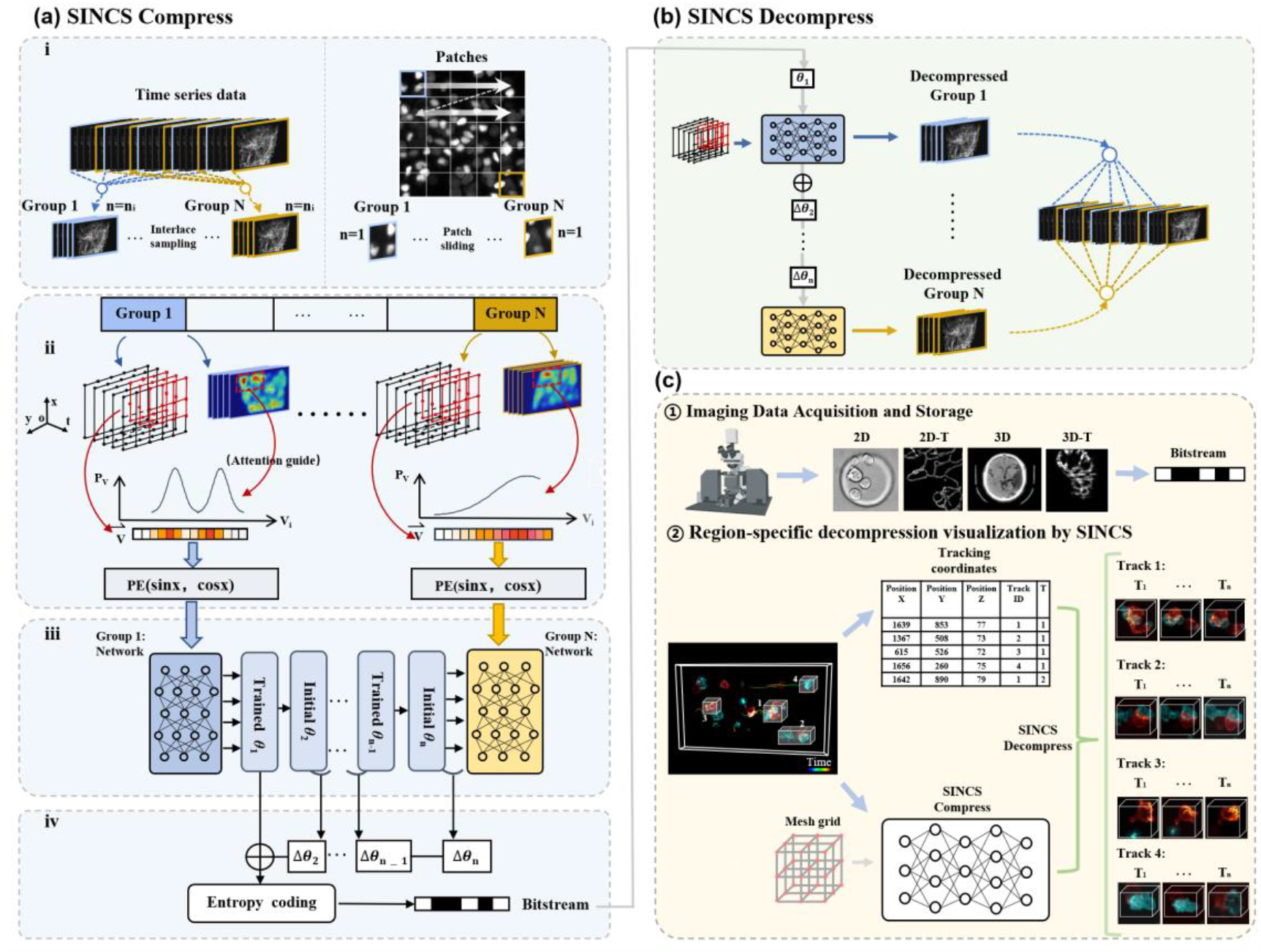
The workflow of SINCS compression and decompression. **(a)** Compression pipeline of SINCS. For a given large-scale biomedical dataset, the pipelie contains: (i) the data is first effectively partitioned into several groups through an adaptive grouping strategy; (ii) saliency mechanism was introduced to realize adaptive compression fitting for biomedical data. This mechanism leverages saliency hot maps (serve as discrete probability distributions for coordinates query) to optimize the compression process,enabling targeted learning of crucial information in the dataset; (iii) Multi-Layer Perceptron (MLP) was constructed as a parameterized mapping function to fit each group data. After saliency-guided sampling, the selected coordinates are first encoded to vectors with high-frequency by positional encoding, and then fed into the MLP,achieving “data-function(weights)” encoding. Subsequently, a weight transfer with trianed parameters *θ*_1_ of 1^st^ group data as the initial parameters and *θ*_2_ of 2^nd^ group data for starting optimization was applied to promote network’s rapid convergence based on the high correlation between groups; (iv) after global fitting of the original data, a higher compression ratio can be further realized using weight-residual entropy coding strategy. Specifically, SINCS encoded residuals by subtracting the network parameters between neighboring networks and applying entropy coding to obtain the encoded initial parameters and residual ones. This step produces a bitstream at last. **(b)** Decompression pipeline of SINCS. In the decompression process, the weight-residual entropy decoding is adopted to convert the bitstream to original initial parameters *θ*_1_ and remaining residual parameters △*θ*_i_. Subsequently, by successively adding the residuals to the initial weight parameters, the original weight parameters are obtained for each network corresponding to each group data. By modeling forward inference and regrouping, the decompressed data can be reordered by the pre-defined grouping strategy. (c) The decompression procedure showing that the region-specific decompression and visualization can be readily achieved in SINCS with flexibility, owing to its coordinate-based representation.

**Fig. 3.**
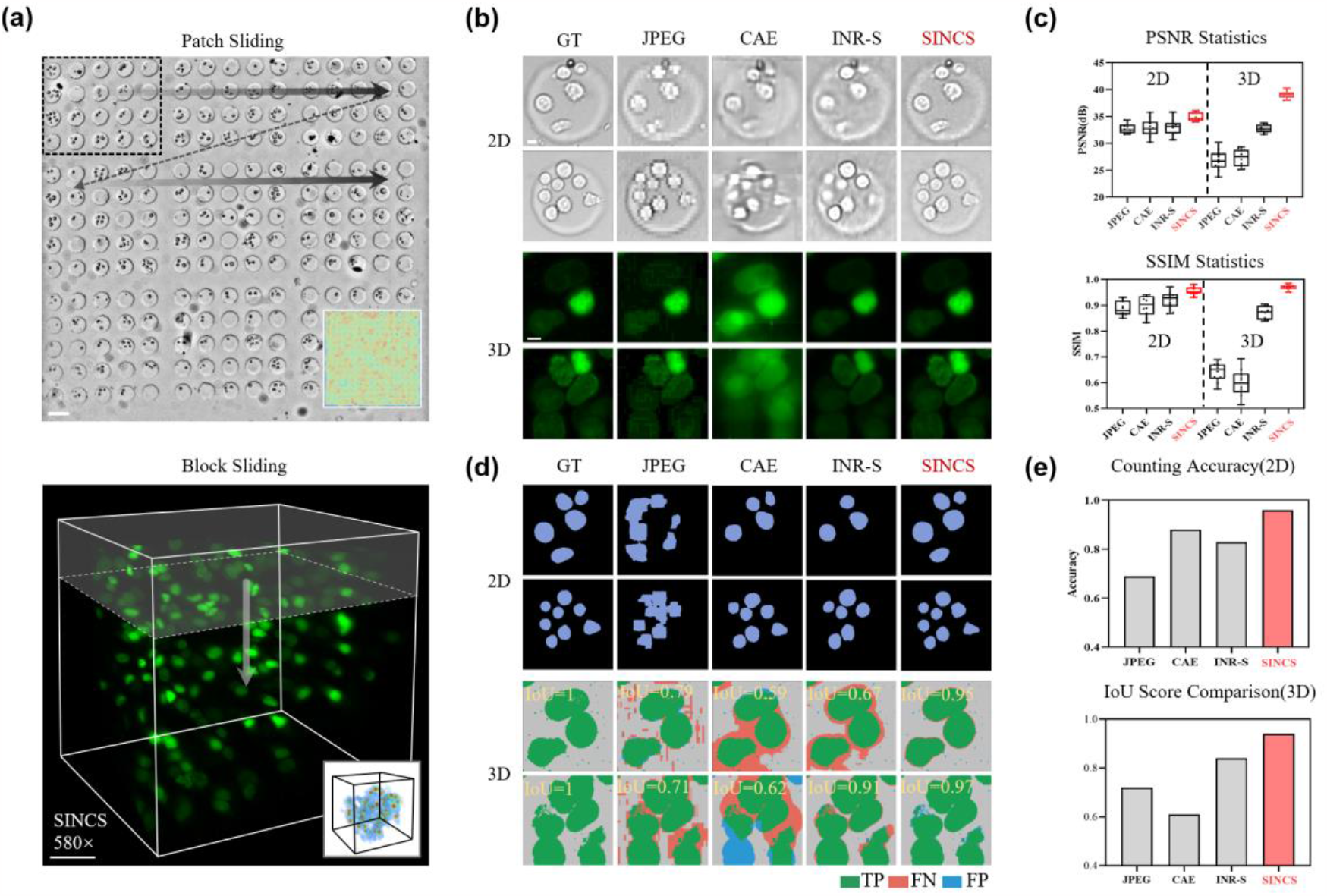
Demonstration of SINCS compression capability on 2D bright field / 3D fluorescence images and performance on downstream tasks. **(a)** Brief illustration of 2D data patch grouping and 3D data block grouping (The respective saliency maps are shown in the bottom right corner). Scale bars from top to bottom: 100 *μm*, 40 *μm*. **(b)** Comparison of bright-field 2D cell data and 3D cell nuclei data (labeled by GFP) reconstructed by different image compression methods. Since JPEG cannot compress 3D data, we convert 3D to 2D data for batch compression, where the data compression ratio is 130× for JPEG (limitation) and 580× for other methods. Scale bar: 5μm. **(c)** Overall performance rating using PSNR and SSIM metrics, to show that SINCS surpass alternative compression approaches in terms of higher structural fidelity. **(d)** Comparative segmentations of 2D cells and intersection over union (IoU) scores of 3D cell nuclei by different compression methods. The IoU scores are used as fidelity metric with higher score indicating higher visual fidelty. TP (True Positive): the correctly reconstructed structures; FP (FalsePositive): the incorrectly hallucinated structures; FN (False Negative): the missing details. Metrics from top to bottom: Counting accuracy, and IoU score comparision. **(e)** Histograms comparing the reconstruction accuracy of different compression methods with using 2D counting accuracy (top) and 3D IoU scores as metric (bottom).

### The principle of SINCS

Biomedical images can be considered as discrete sampling results of continuous spatiotemporal signals. The continuity representation ability of INR is suited for fitting the arbitrary-dimensional signals so that we can map the sampling result to high-dimensional functions, achieving “data-to-function” encoding. However, the large size of biomedical data poses a big challenge for INR network fitting. A simple solution is to increase the parameters of the INR, nevertheless this amplifies the computational complexity and leads to compromised compression efficiency. Based on the locality and repeatability of correlations, we first introduced an interlace group strategy which mitigated this issue through decomposing the massive high-dimensional functions (data) in INR. We used an adaptive grouping strategy to divide the original data into groups (Fig. 2a(i)). For time series data, we specifically made an interlace sampling along time series to divide the original data into N groups, for which both the global motion trend within groups and the high cross-correlation between groups can be retained due to the frame continuity. This temporal grouping strategy helps to reduce temporal redundancy and achieve a higher rate of compression. For patches that lack temporal correlation (obtained by “patch sliding”), the abovementioned grouping strategy can be modified to have n_i_=1 in each group, which is also helpful to the elimination of spatial redundancy. With decomposition, we achieve highly-efficient compression by utilizing multiple simple implicit neural representations (Supplementary Fig. 4).

**Figure 4.**
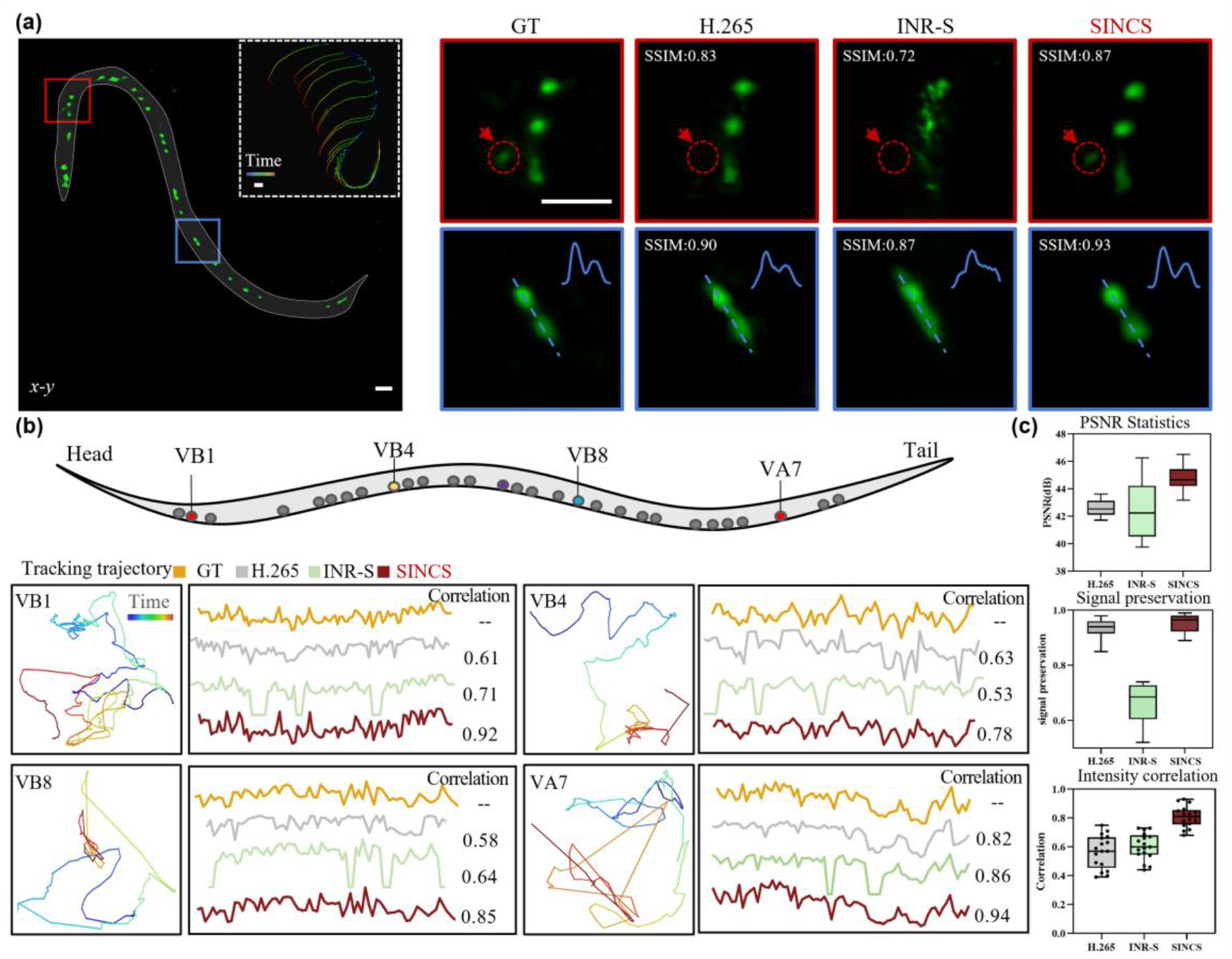
High intensity fidelity SINCS compression on sequential Ca^2+^ images of moving *C. elegans* allowing downstream quantification of neural activities. **(a)** The motor neurons in an entire L4 *C. elegan*s larva reconstructed by different compression approaches (The top right corner shows the *C. elegan*s crawling trend with a time-coded trace). The magnified views of indicated regions show that H.265 and INR-S lose a considerable amount of weak signals, owing to the high signal dynamic range. In sharp contrast, SINCS preserves these weak signals perfectly. Meanwhile, SINCS reconstruction also shows spatial resolution higher than the other two approachs, notably contributing to the resolving of dense signals. The SSIM are used as structral fidelity metric with higher value indicating higher fidelity. Scale bar: 10*μm*. **(b)** Spatio-termporal patterns of 4 motor neurons (VB1, VB4, VB8, VA7) reconstructed by different image compression methods. The neuron tracing trajectories are displayed on the left, indicating the dynamics of neuronal signals in spatial domain. The Ca^2+^ activity curves of corresponding neurons reconstructed by SINCS (red), INR-S (green) and H.265 (gray) approaches are shown on the right, and compared with the ground truth curve plotted by raw data (yellow). The intensity correlations are used as metrics to quantify the intensity fidelity of the reconstructions by diverse methods, with a higher correlation value indicating a higher stability in intensity fidelity. **(c**) Overall performance rating using PSNR (top), signal preservation (middle) and intensity correlation (bottom) metrics, to show that SINCS surpass the H.265 and INR-S in term of both high structural and intensity fidelities.

Then, we designed a saliency-guided sampling to catch structural information of biomedical data. For each pre-defined data group, we proposed a training sampling strategy based on saliency mechanisms (Method section), permitting an adaptive compression that fitted the biomedical data with highdimension structural features. A pre-trained saliency detection network, as illustrated in Fig. 2a(ii), generated saliency hot maps that serve as discrete probability distributions for coordinates query, thus incorporating the structure information into network optimization. This adaptive sampling strategy contributed to contrast improvement in decompressed results, as compared with conventional weighted loss optimization strategy with saliency map (Supplementary Fig. 5).

**Figure 5.**
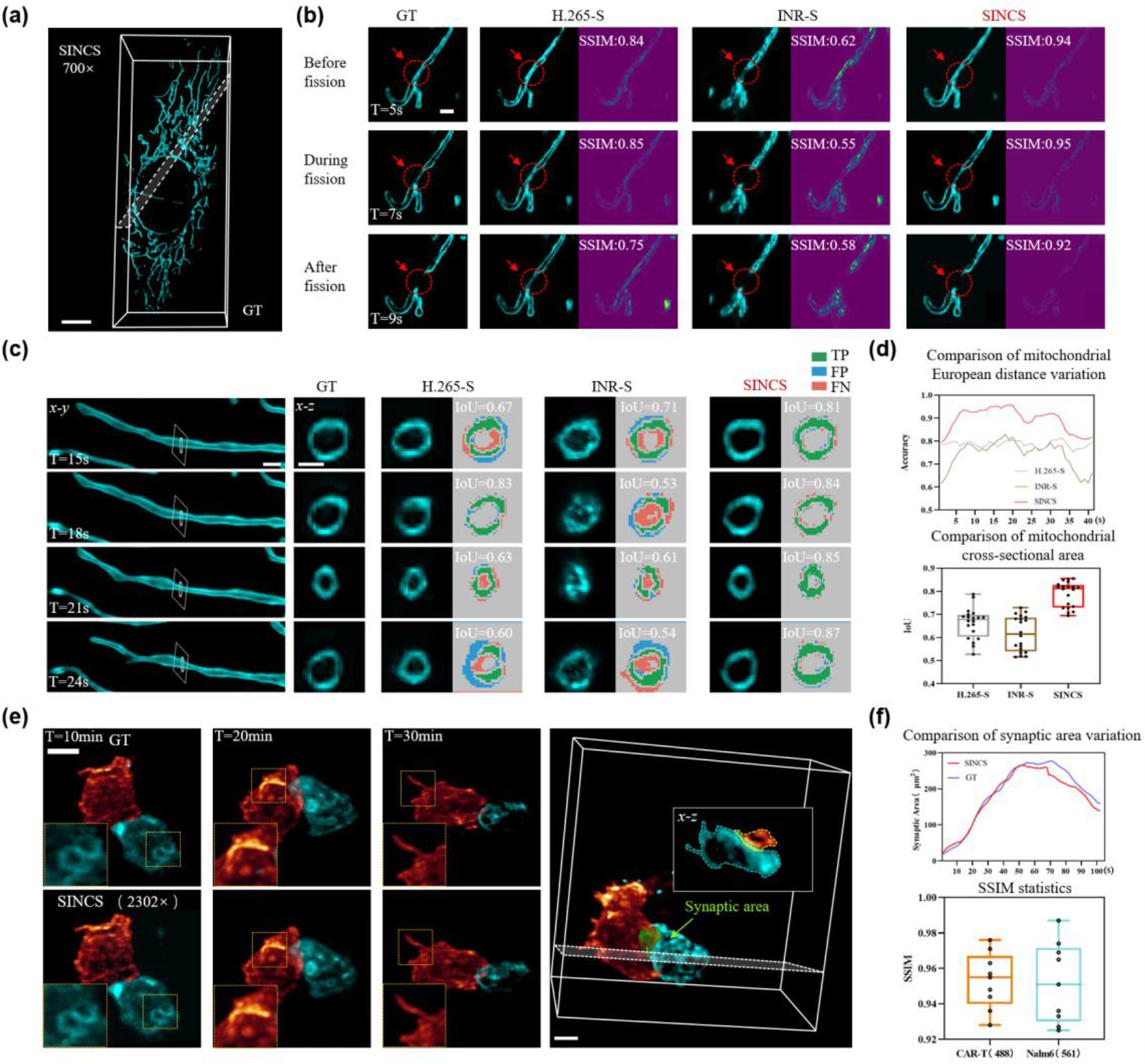
SINCS compression on 4D super-resolution images of mitochondrial dynamics and 5D light-sheet images of CAR-T cell / tumor cell interaction. **(a)** 3D volume renderings of GT (top) and 700× SINCS compression result (bottom) showing the overall high structural similarity by SINCS. Scale bar: 10 *μm*. **(b)** Comparison of mitochondrion fission process reconstructed by H.265-S, INR-S and SINCS.The red arrows indicate the mitochondrion fission site over 4 seconds. As compared to the GT from raw images, only SINCS results are capable of resolving the fine structural changes. The SSIM scores of the reconstructions by three approaches are calculated, with higher value indicating higher fidelity. Scale bar: 1*μm*. **(c)** Comparative results of mitochondrion morphological changes in reconstructed cross section plane. The cross sention plane at different time points demonstrate the mitochondrion contraction and expansion at nanoscale. The IoU scores are used as fidelity metric with higher score indicating higher reconstruction fidelty during the morphological and cross-sectional area changes. TP (True Positive): the correctly reconstructed structures; FP (FalsePositive): the incorrectly hallucinated structures; FN (False Negative): the missing details. Scale bar: 1*μm*. **(d)** Quantitative comparison of reconstruction accuracy at mitochondrial fission site with using European distance as metric(top)and cross-section plane using area as metric (bottom). **(e)** Visual comparison of GT and 2302× SINCS compression result of 5D light-sheet fluorescence microscopy data recording the interactions between CAR-T (labeled by GFP) and Nalm6 tumor cells (labeled by Dsred) in 20 minutes. Scale bar: 5 *μm*. **(f)** Comparison of synaptic area variation and SSIM values between GT and SINCS result.

In the following step, the sampled coordination was converted to corresponding voxels’ value by Multi-Layer Perceptron (MLP), which was a parameterized mapping function to fit data of each group. After saliency-guided sampling, the selected coordinates were encoded to vectors with high-frequency by positional encoding (Fig. 2a(ii)), and then fed into the MLP. During the training process, the parameters of network were updated by the L2 loss computed from the network outputs and original inputs (see Supplementary Note 2, Supplementary Video 1 and Supplementary Table 1 for more training details).

After creating the high correlations between pre-partitioned groups, we designed a weight transfer fine-tuning strategy that adopt sequential fitting of each group data to accelerate network’s convergence. We compressed the data from the first group and obtained the optimized parameters (*θ1*’) for the initial network. Instead of introducing a new model, these optimized parameters served as the starting point for compressing the data from the second group. This meta-learning compression strategy minimizes the distance in INF domain between initial parameters (*θ*_2_) and optimized ones (*θ*_2_*’*) of the 2^nd^ group data (Supplementary Fig. 6). It yielded significantly faster network convergence as compared with direct compression of individual groups. We have demonstrated significant correlation between the weights across various time points in the INF domain, making this weight transfer possible.

With the abovementioned procedure, SINCS successfully achieved rapid compression of arbitrary-dimensional biomedical images into INR weights in a manner of “data-function”. We further increased the compression ratio using weight-residual entropy coding based on the semantic correlation between each group data (Fig. 2a(iv)). SINCS encoded the residuals by subtracting the network parameters between neighboring networks and applying entropy coding to obtain the encoded initial parameters and residual ones for subsequent streamlining storage and transmission processes.

At image decompression stage (Fig. 2b), weight-residual entropy decoding was adopted to convert the highly-zipped bitstream to original initial parameter (*θ*_1_) and remaining residual parameters (△*θ*_i_). After successively adding the residuals to the initial weight parameters, we obtained the original weight parameters of each network for each group data. Through model forward inference and regrouping, the decompressed data can be reordered by the pre-defined grouping strategy. It is noteworthy that with the advantage of coordinate-based representation of INR, region-specific decompression and visualization can be readily achieved for flexible downstream biomedical tasks (Fig. 2c).

### SINCS achieves high-fidelity and high-ratio compression of multidimensional biomedical data

#### High structural fidelity compression for 2D and 3D biomedical images

Bright-field microscopic imaging has been frequently used in biomedical research, generating huge amounts of image data^30^. However, bright-field images typically exhibit low contrast or contain less gradient information, making it more vulnerable to the loss of such gradient structural information during large data compression. Therefore, bright-filed images especially need compression algorithms that can offer both high-fidelity representation and high compression ratios for efficient data storage enabling convenient downstream image-based tasks such as cell segmentation. Here, we applied the SINCS to compress large field-of-view (FOV) 2D imaging data acquired by an inverted light microscope. The field of view of entire two-dimensional plane is 2.67 × 3.99 *mm*^2^, and we divided the plane into 48 patches using patch sliding, as illustrated in Fig. 3a. Each patch was then compressed using the SINCS algorithm guided by saliency map. Considering the large field-of-view information of bright-field imaging, here we take the learnable saliency map to better prioritize the signal information. This learnable saliency map is also applicable to a variety of other modalities of biomedical data, such as CT, MRI, TEM, as shown in Supplementary Fig. 7. To showcase the compression capability, we selected two representative patches for visualization. We further enlarged the regions of interest (ROIs) within these patches in Fig. 3b. Meanwhile, we compared our method with JPEG compression, conventional INR compression and autoencoder-based compression methods (CAE) ^14, 22-24^, demonstrating the superior visual fidelity achieved by our approach. In addition, we applied SINCS to 3D cell data. We partitioned it into multiple blocks using block sliding and rapidly compressed them through weight transfer, as illustrated in Fig. 3a. We visualized the 2D / 3D decompression results of SINCS and those from other compression approaches, showing that our approach offers higher visual fidelity and better preserved cellular details (Fig. 3b). We also quantified the SINCS results using several well-established metrics, showing that it achieved big compression rate of 80 (2D) and 580 (3D), high Peak Signal-to-Noise Ratio (PSNR) of 34.5 (2D) and 38 (3D), and high Structural Similarity Index (SSIM) of 0.94 (2D) and 0.96 (3D). The comparative results in Fig. 3c have shown that SINCS have outperformed the alternative compression approaches in term of these metrics. Furthermore, we conducted image-based cell segmentation using the open-source software “Cellpose”^31^, to validate that the superior compression quality by SINCS also necessarily led to more accurate downstream cell analysis (Fig. 3d). This advantage is further consolidated by quantitatively comparing the Intersection over Union (IoU) scores of the segmented images and the cell counting accuracy (Fig. 3e, Method section). In addition, SINCS with meta-learning also enabled much faster compression as compared to the INR SIREN (INR-S) method that compresses the entire dataset (Supplementary Fig. 8).

### High intensity fidelity SINCS compression on quantitative imaging data of neural activities in moving *C. elegans*

Long-term and high-speed Ca^2+^ imaging of neurons in moving specimens at high spatiotemporal resolution is useful to interrogate the behavior-related neural activities through tracking the Ca^2+^ density change indicated by fluorescence intensity variations^32,33^. Therefore, compression algorithm retaining the signal intensity profile is required in this case, to reduce the data size and also reflect the neural activity state of the samples accurately.

We used SINCS to compress the sequential images of moving *C. elegans* captured by light-field microscopy at a high imaging rate of 100 Hz. we achieved a compression ratio of 1500 folds (From 1.6GB to1.1MB). We further compared the SINCS performance with H.265 and conventional INR on the decompression of Ca^2+^ indicator-labelled motor neurons. The comparative ROIs showed that while H.265 and INR-S lost some weak signals owing to the abrupt intensity variations, SINCS better fit these intensity changes because of the signal enhancement by saliency map (Fig. 4a). It should be also noted that considering the sparsity of neuron signals, we also adopted a conventional threshold-based saliency map to prevent the loss of weak signals (Supplementary Fig. 9). SINCS also outperformed other approaches with showing better resolved dense signals. Then we conducted trajectory tracking of 4 motor neurons (VB1, VB4, VB8, VA7) to investigate their intensity fluctuations during *C. elegans* movement (Fig. 4b). When using the intensity profiles extracted from the raw data as references, we validated that SINCS retained the intensity changes of the dynamic Ca^2+^signals well, surpassing the results from alternative H.265 and INR-S approaches. We further quantified the PSNR, signal preservation rate and intensity correlation based on all the motor neurons (Fig. 4c, Methods section). The inherent mapping function from coordinates to signal values and the adaptability of the saliency map for identifying low-intensity signals together allow SINCS to demonstrate intensity accuracy much higher than other approaches, thereby ensuring authentic representation of dynamic biomedical data and seamless continuation of subsequent tasks (Supplementary Video 2).

### SINCS compression on high-dimensional images of live cells

In long-term live biomedical imaging, a variety of dynamic biological processes, such as blood flow, heartbeat, and cell-cell interactions, occur in four (3D space + time) or even five (3D space + time + spectrum) dimensions and tend to generate tremendous amounts of data which intrinsically need to be compressed. Meanwhile, such types of high-dimensional data are accompanied with complex variations in both temporal and spatial domains, making high-fidelity and high-ratio compression especially challenging and necessary to ensure the downstream tasks being conducted accurately. We applied SINCS to 4D cell super-resolution data which were acquired using our lab-built light-sheet fluorescence microscope (LSFM) with a near isotropic resolution of ∼100 nm. The entire 4D image dataset contains 180 consecutive volumes with totally generating 244 giga voxels (488 gigabytes) to record the 3D dynamics of mitochondrial within a single cell across 3 minutes. SINCS then achieved a 700-fold near lossless compression that drastically reduced the size of the data into 697 megabytes while retained the complex outer membrane morphology. We visualized the decompressed data and the raw data in the same 3D volume rendering (Imaris 9.0) to visually examine the overall high structural fidelity by SINC compression (Fig. 5a). Then we selected three time points of the same small ROI and magnified them to compare the reconstructed details by SINCS, H.265 and INR-S (Fig. 5b). It’s noted that due to the limitation of directly compressing 4D data by H.265, we concatenated all temporal axis data along the axial axis to fit it into 3D format for testing H.265 compression (referring to H.265-S). In visual comparison, SINCS significantly outperformed other approaches, accurately visualizing the transient process of a single mitochondrion fission. The error maps and SSIM metric calculated with using raw image as references further validated that SINCS achieved significantly higher structural fidelity as compared to other approaches. The incomplete mitochondrial fission observed in the results of H.265-S might be from the concatenation of temporal and axial dimensions that led to non-uniform signal distribution and signal residues (Fig. 5b). Meanwhile, since INR-S lacked sufficient fitting ability to learn regions with low signal intensity, it also led to suboptimal structural fidelity (Fig. 5b).

We analyzed the dynamics of a selected mitochondrion at its cross-section plane to further validate the reconstruction fidelity in four dimensions. H.265-S exhibited significant morphological aberrations, likely because of its forced concatenation and fitting along the axial and temporal directions (Fig. 5c, left). In the meantime, INR-S could hardly discern the inner and outer membranes, preventing the subsequent quantitative analyses (Fig. 5c, middle). In contrast, only SINCS achieved smooth morphological changes which are nearly identical with the changes in raw image data. We further quantified the reconstruction accuracy at mitochondrial fission site (metric: European distance) and cross-section plane (metric: area) over time, as shown in the top and bottom of Fig. 5d, respectively. The results verified that the structural fidelity and time signal continuity by SINCS compression were both higher than other approaches. These advances came from our novel interlace grouping strategy that ensured global continuity and inter-frame continuity for accurate compression over time (Supplementary Video 3).

We went deeper with applying SINCS to the light-sheet fluorescence microscopy data recording the interactions between CAR-T and Nalm6 tumor cells, in which the subcellular changes of CAR-T immune synapses and tumor membranes in space, time and spectrum domains together formed a highly complex task for data compression. As we can see in Fig. 5e, while SINCS achieved an impressive compression ratio of 2302 folds, from 1TB to 455.5MB, it also enabled precise cellular morphology reconstruction and thereby accurately reproduced the complete Immunotherapy processes (Fig. 5e). We computed the variation of synaptic areas (Method section) during the interaction between CAR-T cells and Nalm6 T cells to evaluate the temporal compression quality over time. In addition, we calculated the SSIM values within the ROIs across different spectral channels (Fig. 5f), demonstrating that our approach consistently maintains high-fidelity compression performance across various spectral channels and permits reliable data analysis and validation in downstream tasks (Supplementary Video 4).

### Conclusion and Discussion

Both conventional model-based and emerging learning-based approaches show limited performance on the compression of biomedical images that have the features of high dimension, high dynamic range, and often require accurate downstream analysis. SINCS greatly improves the compression of biomedical data in term of performance and applicability by initiating the study on semantic redundancy of biomedical image data in INF domain. After verifying the semantic redundancy in INF domain, we then designed weight transfer optimization strategy and included saliency-guided mechanism adapted to the structural characteristics of multimodal images, making SINCS capable of high-fidelity compression of diverse biomedical data with high compression ratio and improved speed provided.

SINCS applies different grouping strategies for network training and coding based on data types (please refer to Supplementary Note 3, Supplementary Fig. 10 and Supplementary Table 2 and 3 for more details). Then, it generates saliency maps according to the signal distribution of each group of data, which adjust the network training sampling strategy to guide better parameter allocation and achieve adaptive high-fidelity compression. Moreover, since SINCS is an implicit neural function mapping from spatial coordinates to signal values, it can innately incorporate saliency-guided mechanism within it, with significantly-improved fidelity in lateral, axial, and temporal dimensions. Though SINCS is currently not as fast as traditional compression methods yet, its introduction of weight transfer optimization has effectively reduced the model’s training time, as compared with other INR-based compression methods. Also, this reduction in training time will become much more significant, and could be over one order of magnitude when processing increasingly bigger data. In the following decompression process, SINCS only requires simple forward propagation of neural networks, making the decompression speed nearly an-order-of-magnitude faster than H.265, as shown in Supplementary Table 4. It is also worth noting that, for the downstream visualization or quantitative analysis of large-scale biomedical images, multiple transmissions and decompressions may be necessary given the constraints of limited bandwidth. In a lot of practical applications, the high-quality, large-ratio compression as well as high-speed decompression by SINCS makes it outstanding from the alternative approaches.

We validate the capabilities of SINCS on several types of biomedical images, especially on 4D super-resolution microscopy data of live cells whereas high-resolution, high-fidelity, efficient compression are all required. When facing these challenging data with large size, dynamic structures and high resolution, SINCS approach notably outperforms traditional H.265 and INR-based method, rendering itself a powerful and versatile data compression and transmission tool for diverse biomedical applications. Moreover, to specifically address the loss issue for certain types of medical datasets, SINCS can achieve true lossless compression by further incorporating image residuals, as demonstrated in Supplementary Fig. 11. We envision that the INR-based compression could be more versatile with obtaining image priors of diverse samples through meta learning.^34^ We also anticipate the further reduction of compression training time by continuously optimizing the network design strategies.^35^

## Methods

### Network optimization strategy based on saliency mechanism

We introduce a saliency mechanism with learnable or hard saliency maps to guide the adaptive allocation of parameters in INR, bridging the gap between INR and data and achieving improved data compression fidelity, as shown in Supplementary Fig. 12. Considering the characteristics of the signal distribution, we have two different saliency maps to cope with signals of different distribution types. Specifically, for data with multiple ROIs, dense signal distributions, and a demand for high structural detail, we use Gradient-weight Class Activation Mapping (Grad-CAM) ^36^ and Multi-Structure Region of Interest (MS-ROI) ^37^ techniques to create learnable saliency maps for the corresponding data, assigning probability values to each grid coordinate point, reflecting its proportionate importance in the data. In the process of generating saliency maps, we first apply a convolutional layer to the original data, denoted as x (with a size of *a×b*), using a convolution filter of size *n×n*. The convolution operation is represented by Equation 1, where W represents the learned filter.

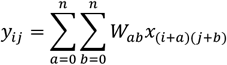

In practice, multiple filters within each layer are learned in parallel, resulting in a three-dimensional feature map as the output of the convolutional layer, where the depth represents the number of filters. Subsequently, by utilizing the learned weights between the predicted class c and the feature map d, we train the Class Activation Maps (CAM) model to obtain saliency maps that capture the significance of the predicted class distribution.

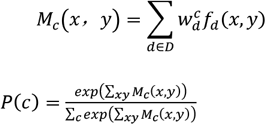

where 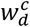 is the learned weight of class *c* for feature map *d*. Training for CAM minimizes the cross entropy between objects’ true probability distribution over classes (all mass given to the true class) and the predicted distribution. The probability *P*_*c*_ represents the likelihood of selecting the corresponding coordinate point for each training iteration. Besides, for sparse signals with faint intensity and lack of structural details, since the subsequent task analysis focuses only on their spatial location or intensity information, we use hard saliency maps based on threshold divisions to prevent the loss of weak signals. After obtaining the saliency maps, the coordinate vectors *V* guided by saliency maps are further mapped to a high-dimensional embedding space using position encoding, enhancing perceptual quality. Formally, the encoding function employed in our approach is as follows:

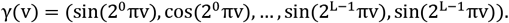

Here *γ* is the mapping of the original coordinate vector *V* from *R* to *R*^2*L*^ and *L* is the number of frequencies used.

### TSNE clustering dimensionality reduction and correlation analysis

For correlation analysis in implicit neural function domain and spatial domain, we adopt Average Wilson Coefficient (AWC) to compare the correlation between two discrete distributions. WC can be computed from the Kolmogorov Smirnov (KS) test formula as follows:

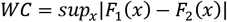

where *F*_1_*(x)* and *F*_2_*(x)* are the Empirical Cumulative Distribution Function (ECDF) of the two distributions, respectively. In this paper, the distributions were defined as the one-dimensional vectors reshaped from the compressed network weights and the original images, respectively. The network parameter values and image pixel values are all normalized to 0 to 1. The smaller value of AWC indicates the higher correlation in the parameter value distributions between the two samples.

To interrogate the statistic correlation among sample, we use t-SNE^29^ dimensionality reduction and clustering to get the data distribution, and we adapt Rayleigh entropy to quantify the degree of clustering, which is calculated as follows:

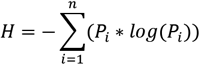

where *P*_*i*_ is the probability that the sample point belongs to a category in the clustering result. A lower Raleigh entropy value indicates that the data distributions are more concentrated in the clustering result.

### Evaluation Metrics

PSNR, SSIM and IoU scores were used in our work to evaluate the compression quality of the decompressed data with respect to the original data. (PSNR, SSIM and IoU scores are all based on single channel images.) Denoting 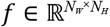 as the decompressed data, and 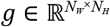 as the original data,PSNR and SSIM values were calculated using the following equations (Take a 2D image as an example):

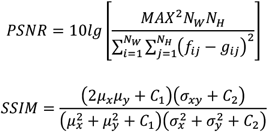

Here *MAX* stands for the dynamic range of the original data. *M*_*x*_, *μ*_*y*_ and *σ*_*x*_, *σ*_*y*_ are the mean value and the standard deviation of the original data and decompressed data, respectively. And *C*_*1*_ and *C*_*2*_ are constants to avoid a zero denominator. It can be deduced that compression quality is better when the SSIM is closer to 1.

And the Intersection over Union (IoU) scores:

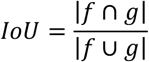

Here |*f* ∩ *g*| represents the size of the intersection between the decompressed data region and the original data region. |*f* ∪ *g*| represents the size of the union between the decompressed data region and the original data region. IoU takes values between 0 and 1, with values closer to 1 indicating a higher degree of overlap between the decompressed data and the original data.

### Quantitative intensity correlation analysis of worms

We performed semi-automatic tracking of motion and intensity fluctuations in each neuron of the GT using the TrackMate Fiji Plugin^38^. Neurons in each volume were automatically detected by applying a circular ROI through a Difference of Gaussian (DoG) detector and then tracked using a Kalman filter. If the automatic tracking failed due to rapid neuronal movement, manual correction of missing detections and tracking errors was required. After tracking neurons in the GT dataset, we export the corresponding neuron’s position coordinates to Excel. Subsequently, leveraging a custom localization algorithm, we perform localization on results under different compression methods. This process entails extracting the average intensity value *F*_*i*_ of all pixels within the ROI surrounding the neuron’s coordinates, effectively representing the fluorescence intensity of that neuron. Finally, employing the same approach outlined above, we generate intensity change curves *L*_*T*_ for neurons under different methods and compare them to the ground truth intensity change curves *L*_*R*_.The intensity correlation is calculated as *correlation* =*L*_*T*_/*L*_*R*_, representing the degree of intensity correlation.

### Cell contact area analysis

We designed an algorithm to quantitatively analyze the contact area between immune cells and cancer cells during their interaction. Initially, a deep learning-based segmentation network^39^ is employed to segment immune cells and target cells. Subsequently, based on the segmentation results, the image is divided into four regions: immune cells, cancer cells, background, and the boundary region. Finally, in the boundary region, distance transformation and watershed algorithms are used to obtain the segmented results of the contact area between immune cells and cancer cells. The segmentation results ensure a single-pixel thickness, enabling the conversion of pixel count into the contact area.

### Sample preparation

Transgenic zebrafish lines *Tg(gata 1a:dsRed;cmlc2:gfp)* was used in our experiments. Embryonic fish were maintained at 3-4days post-fertilization in standard E3 medium, which was supplemented with extra 1-phenyl 2-thiourea (Sigma Aldrich) to inhibit melanogenesis. Then, the larvae were anesthetized with tricaine (3-aminobenzoic acid ethyl ester, Sigma Aldrich) and immobilized in 1% low-melting-point agarose inside a fluorinated ethylene propylene tube for further imaging.

For 2D cell data, cell cultures were prepared using homemade microchips. T-cell medium was used to replace the sterile water, and 500μL of the medium was kept in the confocal dish to submerge the chip. Then, 60μL of CAR-T cells at a density of 1×10^6^/mL was taken and dropped in. We waited for 10 minutes to allow the cells to fall into the chamber. Subsequently, an equal amount of target cells was taken, and the above operation was repeated.

MCF-7 cell line that expresses GFP endogenously was used in 3D cell nuclei data compression experiment, MCF-7 cells were grown in Dulbecco’s modified eagle medium (DMEM), which were supplemented with 10% fetal bovine serum (FBS) and 1% penicillin-streptomycin. Once the cells had grown to 80–90% confluence, they were harvested by 0.25% Trypsin-EDTA treatment and resuspended in the complete medium to a suspension volume of 1× 10^6^ cells/mL.

The strain ZM9128 *hpIs595[Pacr-2(s)::GcaMP6(f)::wCherry]*, expressing GcaMP6f in A- and B-class motor neurons, was used to detect neuronal activity in the moving worm. The *C. elegans* were cultured on standard nematode growth medium plates seeded with OP50 and maintained at 22°C incubators until the L4 stage.

To label microtubules in live U2OS cells, we followed a previously described protocol^40^, in which the cells were coincubated with 4 μM PV-1 and 5 μM Tubulin-Atto 488 at 37 °C for 1 h, then the cells were washed three times with culture medium (warmed to 37 °C) and cultured at 37 °C for another 1 h. Finally, the medium was replaced with phenol red free McCoy’s 5A medium and imaged via DR–SPIM. For labeling mitochondria in fixed cells, U2OS cells were first transfected with Tomm20-EGFP (mito OM) or Cox4-EGFP (mitochondrial matrix) using Lipofectamine LTX according to the standard protocol and cultured at 37 °C with 5% CO_2_ for an additional 24 h. Before imaging, the cells were fixed with 2% glutaraldehyde for 20 min.

For multi-channel 4D biomedical data compression experiment, Acute B-lymphocytic leukemia cell line Nalm6 were cultured in RPMI 1640 medium (Gibco, Grand Island, NY, USA) containing 10% fetal bovine serum (FBS; Gibco, Grand Island, NY, USA). The lentivirus packaging cell line LentiX™293T was cultured in DMEM medium (Gibco, Grand Island, NY, USA) supplemented with 10% FBS. CAR-T cells were pretreated with 50 nM dasatinib (Selleck, Shanghai, China) for 24 h. Due to the reversible effect of dasatinib, 50 nM dasatinib was also added to all subsequent staining, imaging, and other experimental solutions.To label microtubules, CAR-T cells were stained with the SiR-tubulin probe (SpiroChrome, Switzerland) at 2 μM final concentration and incubating for 1 h in a humidified 5% CO2 incubator at 37 °C. The cells were then washed twice with warm phosphate buffer saline (PBS) and resuspended with imaging solution, consisting of the phenol red-free 1640 medium (Gibco, Grand Island, NY, USA) supplemented 10% FBS, 25 mM HEPES (Gibco, Grand Island, NY, USA), 100 U/ml penicillin and streptomycin (Gibco, Grand Island, NY, USA), and 1 μM SYTOX™ Blue stain (Invitrogen, Waltham, MA, USA).

### Data acquisition

For moving *C. elegans* data compression experiment, the calcium signals of worms were captured using a customed-built light-field microscope (LFM)^41^. A water immersion objective (LUMPlanFLN 40×/0.8w, Olympus) was used to collect the epifluorescence signals from samples with scientific camera sensor (Flash 4.0 V2, Hamamatsu). The captured light-field sequences were subsequently reconstructed by the trained VCD model^41^ to yield the 3D videos of the calcium signals in moving worms. The max intensity projection of the 3D reconstructions was then used for the compression.

For 2D and multi-channel 4D biomedical data compression experiment, the multi-channel fluorescently labeled Car-T cell images and bright-field cell images were captured using a customed single objective light sheet microscopy compatible with both fluorescence and bright-field capabilities, based on the IX83(Olympus) framework. The primary optical elements in this configuration include the following: Objective O1 (UPLSAPO 60×/1.35, silicone, Olympus), Objective O2 (UPLXAPO 40×/0.95, air, Olympus), and Objective O3 (AMS-AGY v2.0). This system attains a spatial resolution of 0.35 × 0.35 × 1μm (with axial resolution enhancement through post-processing). For bright-field cell data, illumination was provided by LED light sources, and image acquisition was performed using an Andor camera with an exposure time of 20ms. For multi-channel fluorescently labeled Car-T cell data, excitation was conducted using lasers at 488 and 561 nm, with the 488-channel being captured by a Hamamatsu camera with a 200-ms exposure time, and the 561-channel requiring a 2000-ms exposure time. The color filter for channel 488 is MF525-39, and the color filter for channel 561 is FBH600-40.

For static 3D biomedical data experiment, the 3D cell nuclei data was collected by a customed dual-objective light sheet microscopy. The primary optical elements in this configuration include the following: illumination objective (Mitutoyo Plan Apo Infinity Corrected Long WD Objective 20×/0.42, air), and detection objective (UPLFLN20XPH 20×/0.5, Olympus). This system attains a spatial resolution of 0.325 × 0.325 × 0.5μm. The excitation source of the system is a laser with a wavelength of 405nm, while the sCMOS camera (Orca Flash4.0 v.3, Hamamatsu) acquires data with an exposure time of 20ms.

For 4D biomedical data experiment, 4D cell super-resolution data was collected by a customed dual-objective light sheet microscopy, followed by post-processing image enhancement using an ID neural network^42^. The fluorescence signals generated within the specimen were collected by a detection objective (LUMFLN 60×/1.1 W, Olympus). The resolution of the system is 97×97×450nm. The sCMOS camera (Orca Flash4.0 v.3, Hamamatsu) exposure was precisely triggered with a minimal 2-ms delay to effectively reduce motion blur, and the camera recorded the plane images at a rate of up to 1,000 fps.

The CT, MRI, and TEM data used in our experiments were taken from publicly available datasets. The CT images from the publicly available dataset on http://headctstudy.qure.ai/dataset,MRI images from the publicly available dataset on http://adni.loni.usc.edu/data-samples/access-data/, and TEM images from the Virus Image Dataset (aggle.com).

## Data availability

The datasets generated and analyzed in this study are available from the corresponding authors upon reasonable request.

## Code Availability

The data and code that support the findings of this study are available from the corresponding author upon reasonable request.

## Acknowledgements

We are grateful to Dr. Zhang Meng and Dr. Yuhui Zhang for providing us the fluorescent cell samples. This work was supported by the funding from National Key Research and Development Program of China (2022YFC3401102). National Natural Science Foundation of China (T2225014, 21927802).

## Author contributions

P.F., Y.M. and B.L. conceived the idea. P.F., and B.L. oversaw the project. Y.Z., S.G., L.Z., Y.Z., J.W. and Z.W. developed the optical setups and acquired the experimental images. Y.M., C.Y., X.Y., J.L. and Z.W. developed the programs. Y.M. and C.Y. processed the images. Y.M., C.Y., B.L. and P.F. analyzed the data and wrote the paper.

## Competing interests

The authors declare no conflicts of interest.

## Notes

### Competing Interest Statement

The authors have declared no competing interest.

### Summary of Updates

We add more experiments as well as verification of the principles, and provide detailed supplementary materials.

