## Supplementary materials for "Semantic redundancy-aware implicit neural compression for multidimensional biomedical image data"

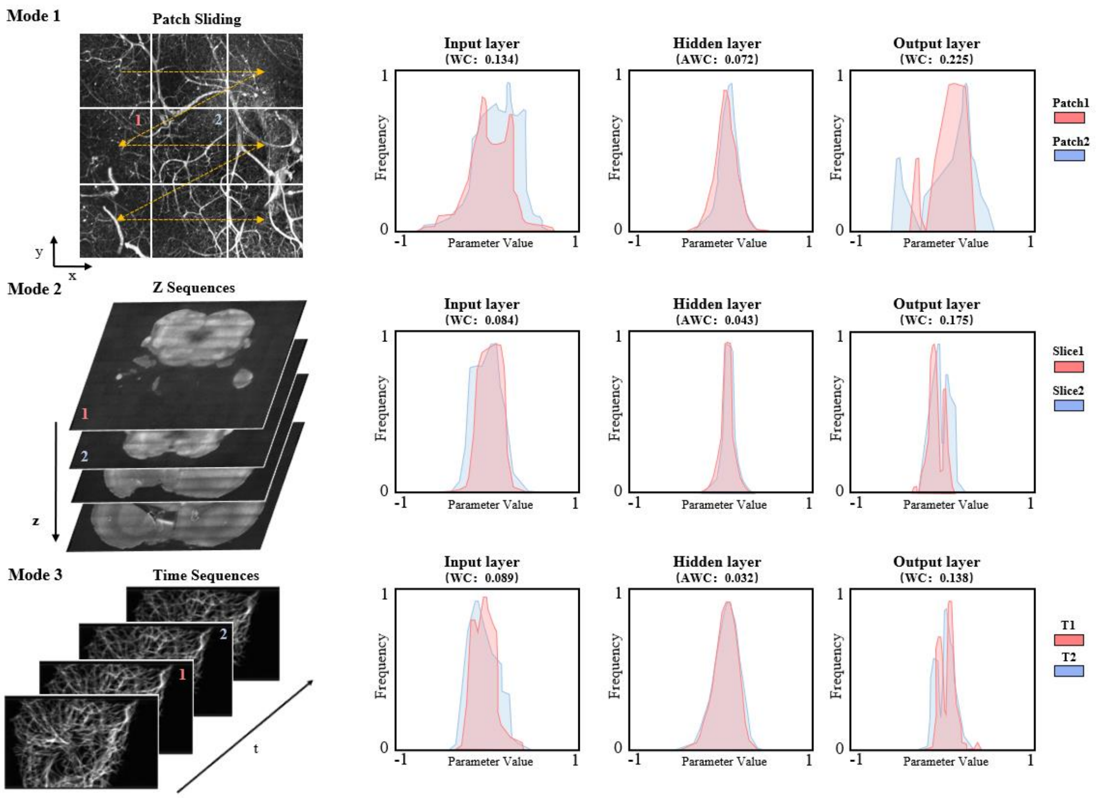


**Supplementary Figure 1. Comparison of distribution of parameters at different layers of the network within the implicit neural function domain in three modes.** In each mode, the semantic correlations of different network layers in the implicit neural function domain are compared through calculating their parameter histograms. The Average Wilson Coefficient (AWC) value is used as correlation metric with lower value indicating higher correlation.From left to right: Input layer, Hidden layer, and Output layer.

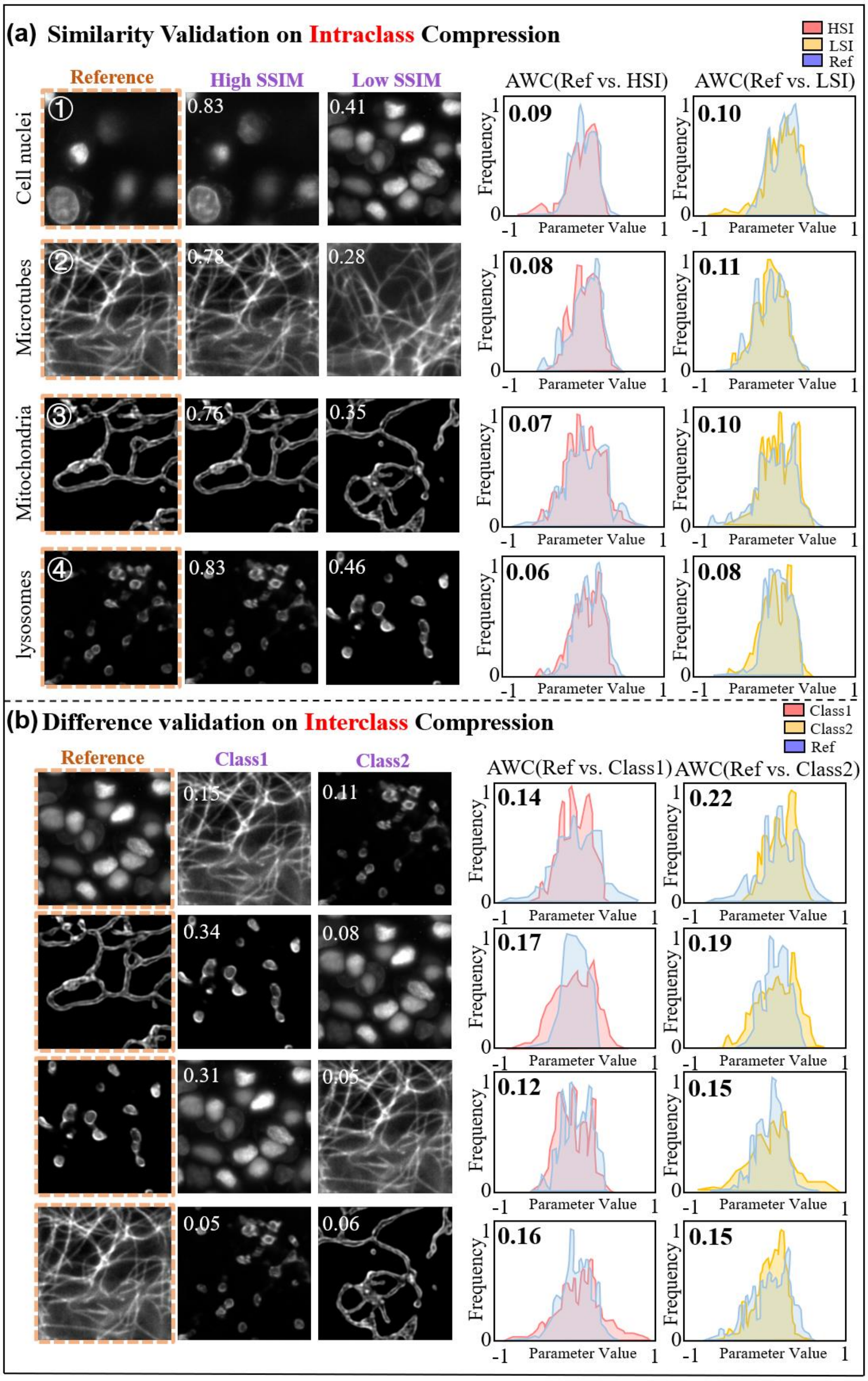


**Supplementary Figure 2. Comparison of intra- and inter-type disparities among diverse types of intracellular organelles (cell nuclei, microtubes, mitochondria, lysosomes) in both the spatial domain and the implicit neural function domain.** **(a)** Quantitative comparison of parameter distribution disparities within the implicit neural function domain for the same type of biomedical images but with different similarity. **(b)** Quantitative comparison of parameter distribution disparities within the implicit neural function domain for different types of biomedical samples.

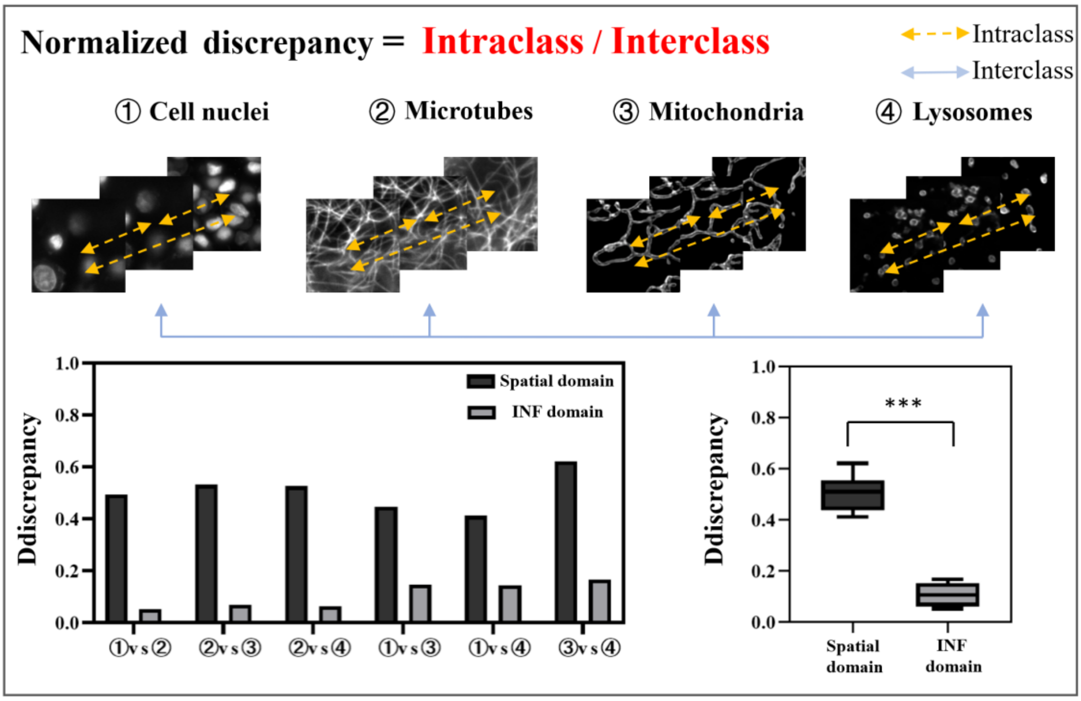


**Supplementary Figure 3. Statistical comparison of intra- and inter-type disparities among diverse types of intracellular organelles (cell nuclei, microtubes, mitochondria, lysosomes) in both the spatial domain and the implicit neural function domain.** The discrepancies among the structures in spatial and INF domains are quantified by the SSIM (black) and AWC (gray) metrics, respectively, with using the equation described in **Supplementary Note. 1** (bottom-left). It's quite obvious that the variation of discrepancy in INF domain is significantly smaller than that in spatial domain.

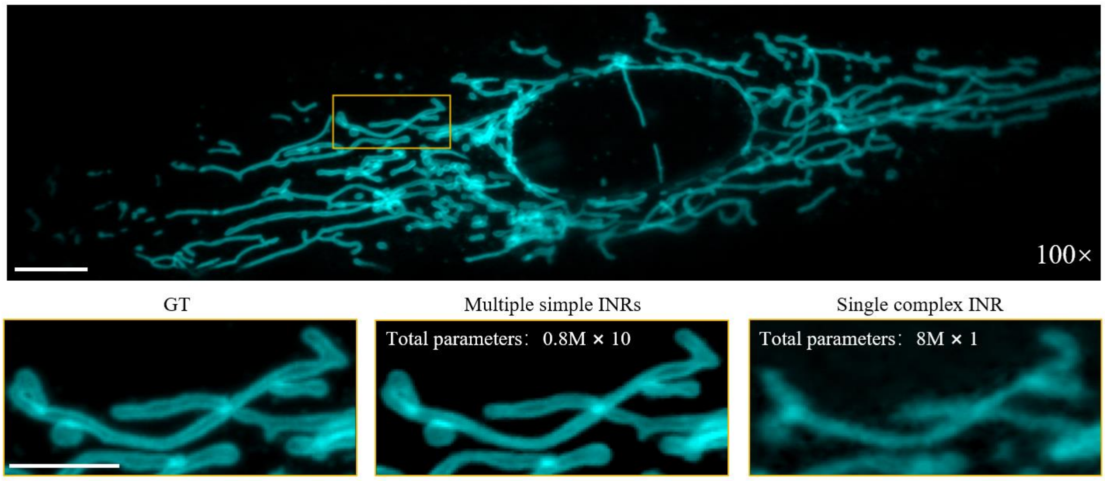


**Supplementary Figure 4. Comparison of compression results using multiple simple implicit neural representations and single complex implicit neural representation.** Visual comparison of the compression results of mitochondrial images with multiple simple INRs and single complex INR at an identical compression ratio of 100×. Bottom: Close-up views of the ROI annotated in the original image. Scale bars: 10 $\mu m$ (top), 5 $\mu m$ (bottom). The images of mitochondrial are captured by a light-sheet microscope using 60×/1.1 NA detection objective.

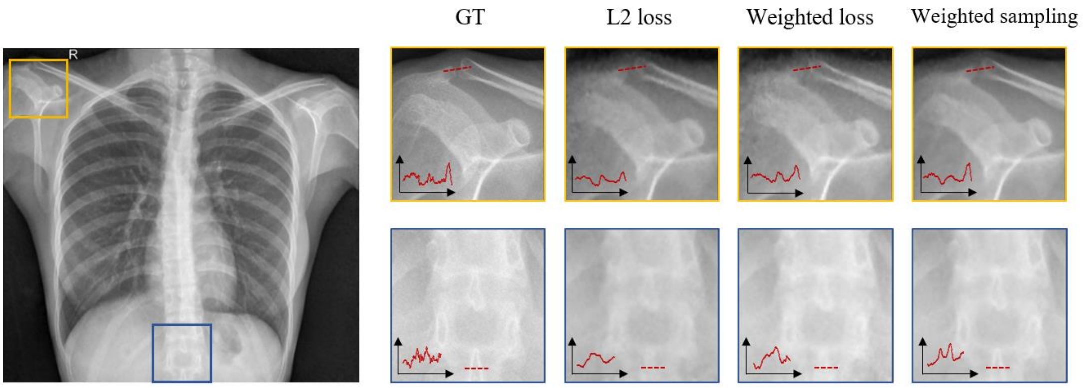


**Supplementary Figure 5. Comparison of INR-based compression with / without learnable saliency map guiding the network sampling and loss function.** The compression results of the model adopting the standard L2 norm loss function is affected by random uniform sampling optimization, resulting in low fidelity in the signal regions; The model utilizing the saliency-guided L2 loss function (Weighted loss[1]) improves compression quality and contrast of the signal regions. Specifically, we introduce the importance probability coefficients of the saliency map as corresponding weights into the optimization of the loss function calculations; The compression results of the model employing saliency-guided sampling (Weighted sampling) further improve the compression quality owing to the optimized training for signal regions. (Compression results from left to right: GT, INR compression employing the L2 norm loss function, INR compression utilizing a saliency-guided L2 loss function, and SINCS employing saliency-guided sampling combined with the L2 norm loss function.)

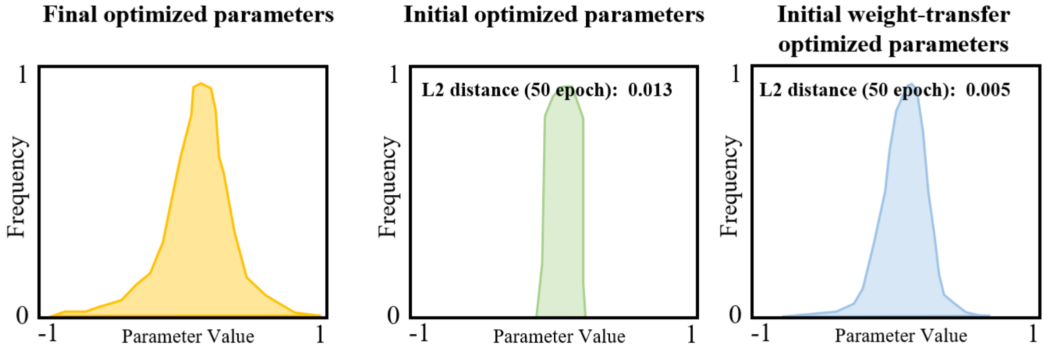


**Supplementary Figure 6. Comparison of the L2 distance in implicit neural function domain between initial optimized parameters (with and without weight-transfer strategy) and final optimized parameters.** The correlation in the implicit neural function domain between the initial optimized parameters (50 epoch) and final optimized parameters (10000 epoch) are compared through calculating their L2 distances. The L2 distance are used as correlation metric with lower value indicating higher correlation.

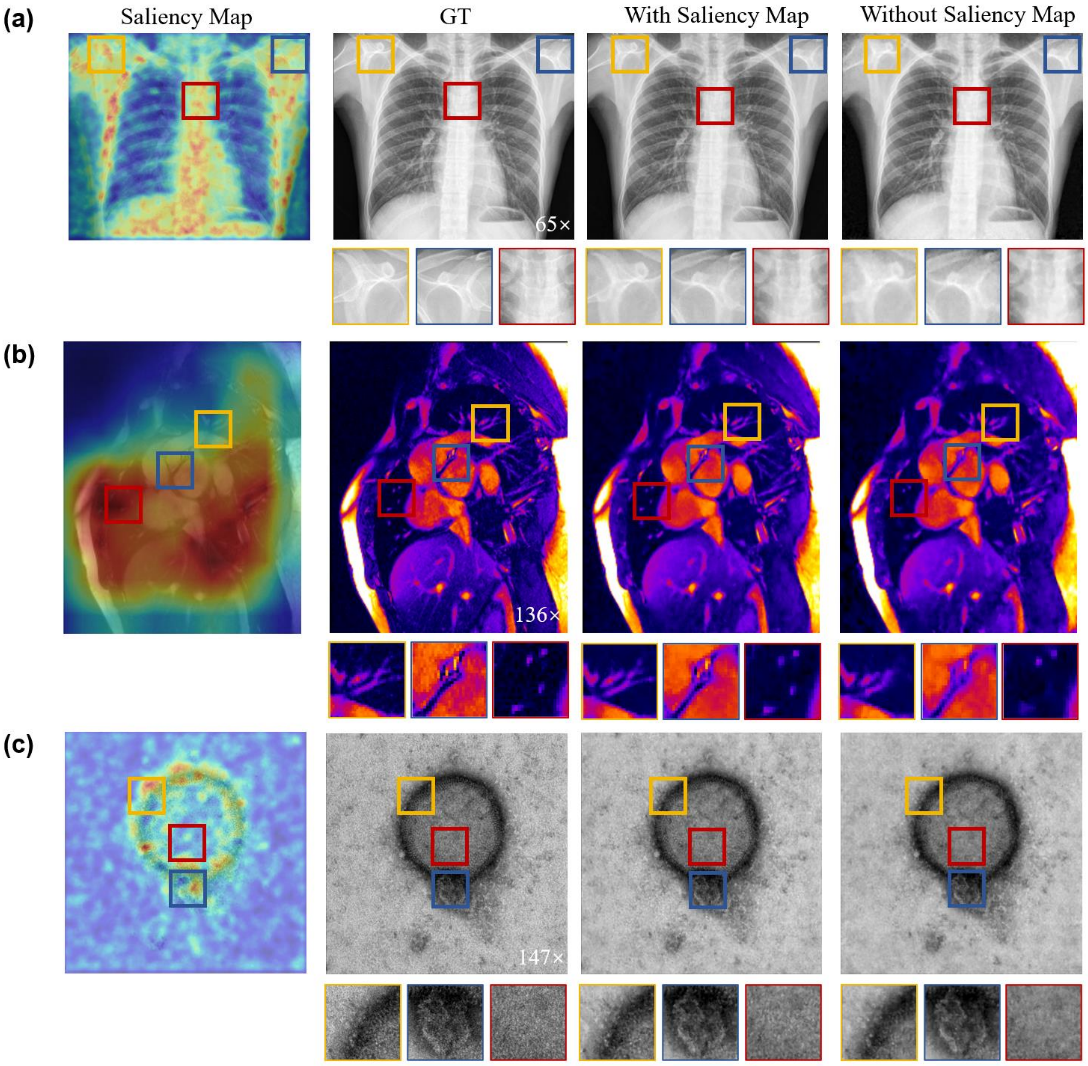


**Supplementary Figure 7. Comparison of compression results with and without the guidance of learnable saliency maps. (a)** Visual comparison of the compression results of CT image of human chest with and without guidance from saliency map at a compression ratio of 65×. Bottom: Close-up views of ROIs annotated in the original images. **(b)** Visual comparison of the compression results of MRI image of human heart with and without guidance from saliency map at a compression ratio of 136×. Bottom: Close-up views of ROIs annotated in the original images. **(c)** Visual comparison of the compression results of TEM image of virus with and without guidance from saliency map at a compression ratio of 147×. Bottom: Close-up views of ROIs annotated in the original images. (CT images from the publicly available dataset on http://headctstudy.qure.ai/dataset, MRI images from the publicly available dataset on https://data.mendeley.com/datasets/, and TEM images from the Virus Image Dataset (kaggle.com)).

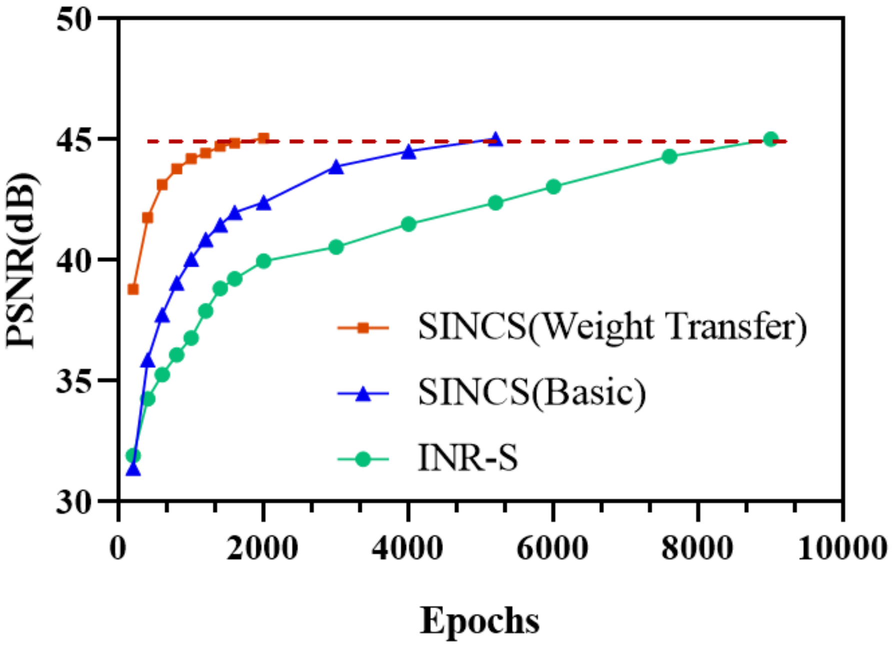


**Supplementary Figure 8. The required training iterations of SINCS with weight transfer strategy, SINCS without weight transfer strategy, and INR-S compression.** With the weight-transferred fine-tuning strategy, SINCS is able to start network optimization with initial parameters much closer to the final optimization parameters. Therefore, it significantly improves the network training speed and reduces the compression time. In this experiment, we compressed 10 groups of fluorescently labeled CART cell data and took their average to present the results.

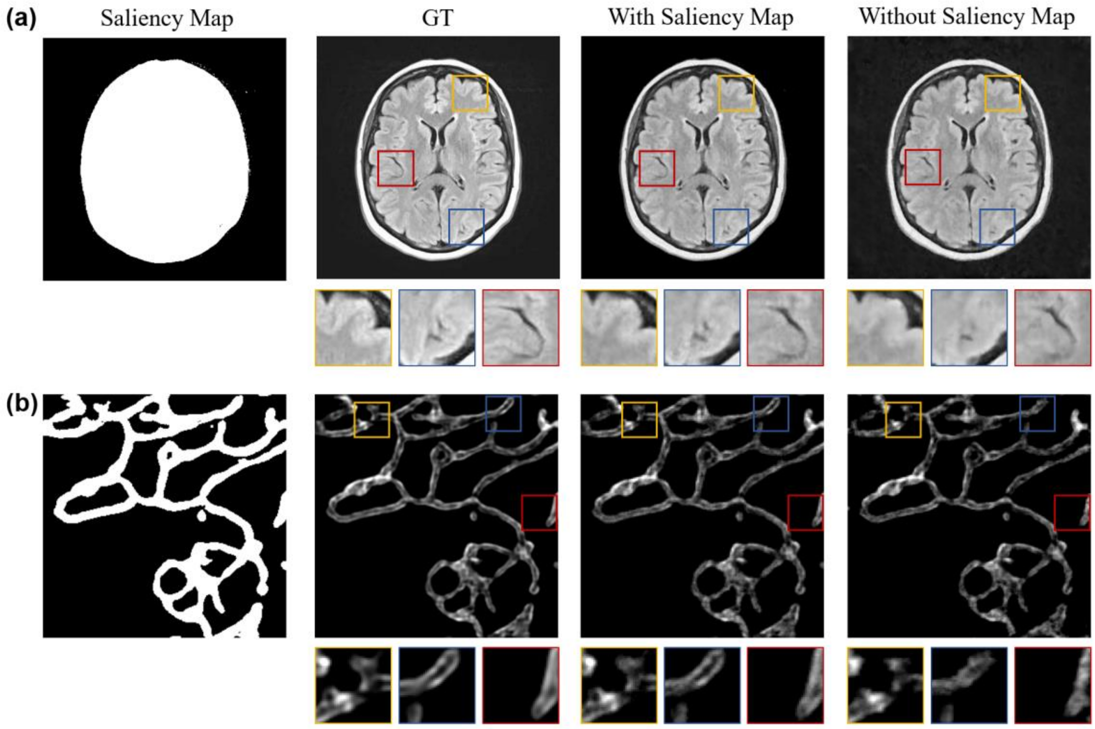


**Supplementary Figure 9. Comparison of compression results with and without the guidance of hard saliency maps. (a)**Visual comparison of the compression results of MRI image with and without guidance from saliency map. Bottom: Close-up views of ROIs annotated in the original images. **(b)** Visual comparison of the compression results of light-sheet microscopy image with and without guidance from saliency map. Bottom: Close-up views of ROIs annotated in the original images. (MRI images from the publicly available dataset on https://data.mendeley.com/datasets/).

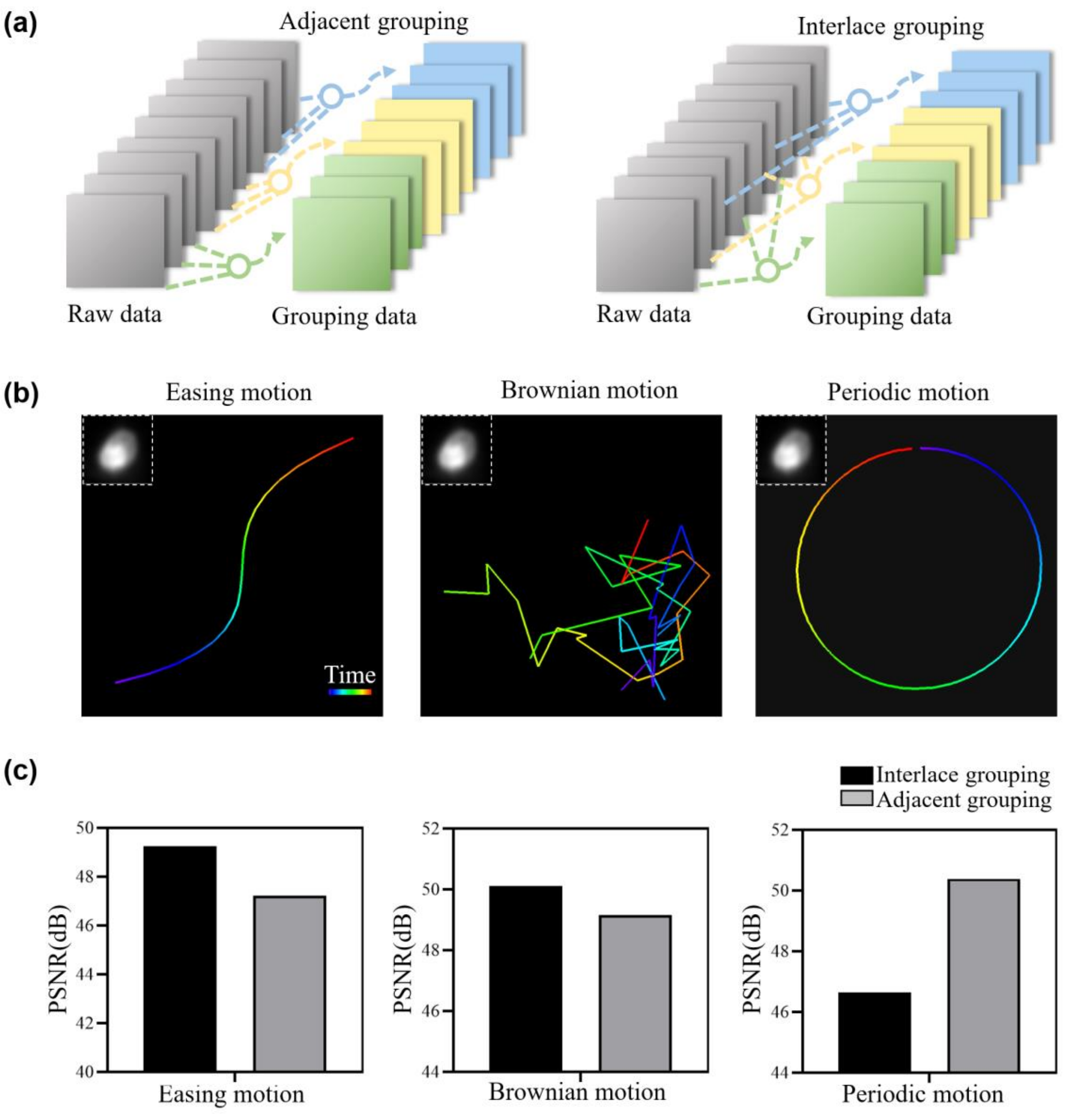


**Supplementary Figure 10. Compressing simulated data with three different motion types using two different grouping strategies. (a)** Schematic of two different grouping strategies (Adjacent grouping and Interlace grouping). **(b)** Simulated cell data for three different motion types (Easing, Brownian, Periodic). **(c)** PSNR statistics of simulated cell data of different motion types in two grouping strategies.

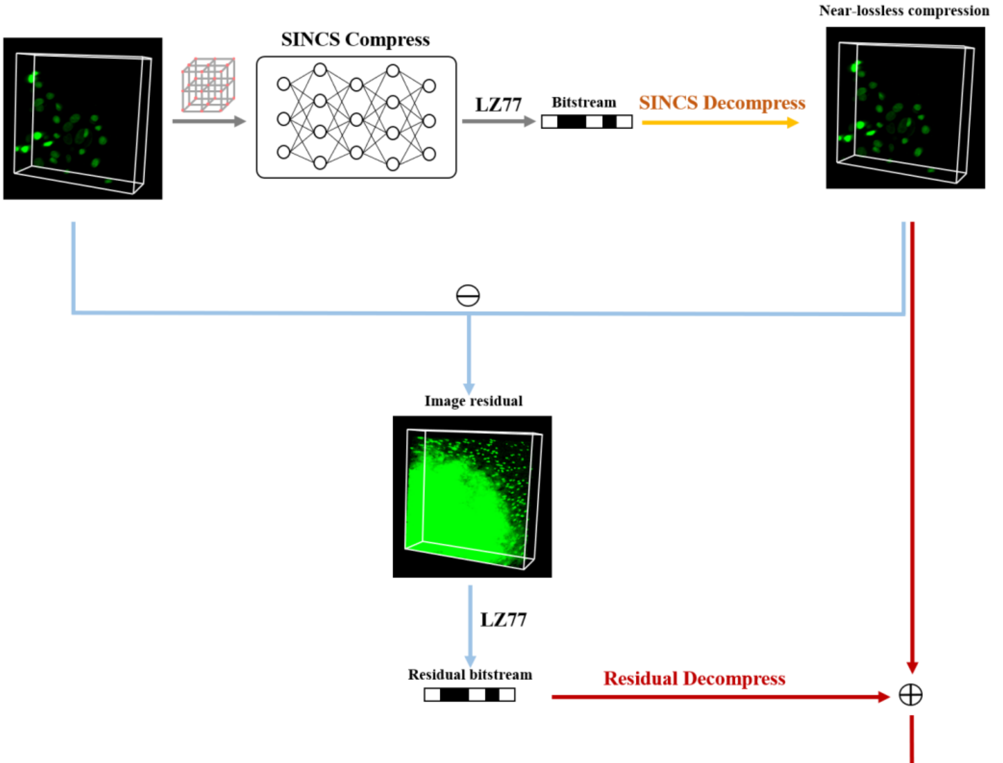


| Methods | Raw data size | Data size after compression |
|---|---|---|
| LZ77 (Lossless compression) | 203MB | 68.4MB |
| SINCS (Near-lossless compression) | | 368KB |
| SINCS (Lossless compression) | | 59.4MB |

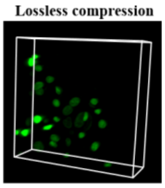


**Supplementary Figure 11. The brief schematic of the two types of SINCS compression (near-lossless and lossless).** To 100% restore the signals for certain types of medical datasets, SINCS can also achieve true lossless compression by further incorporating image residuals. The table compares the compression ratio between near-lossless SINCS compression, true lossless SINCS compression, and classic Lempel-Ziv 1977 (LZ77) method[2].

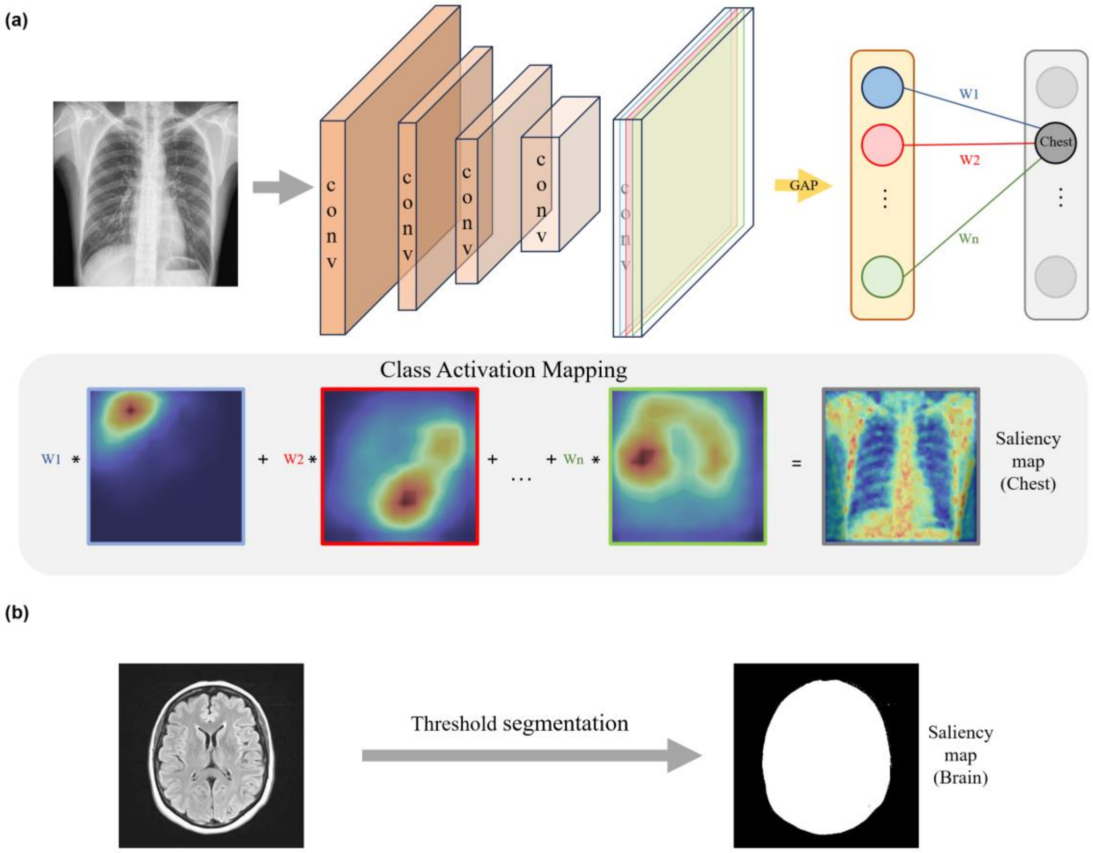


**Supplementary Figure 12. Schematic diagrams showing the generation of two types of saliency maps. (a)** The schematic diagram of the learnable saliency map generation. ("Conv" denotes the convolutional layer, and "GAP" denotes Global Average Pooling. The weights '$W_i$' for each channel in the feature map are obtained through global average pooling. Subsequently, a linear combination of these weights is computed to yield the learnable saliency map. **(b)** The schematic diagram of the hard saliency map generation simply using the threshold segmentation.

### Supplementary Note 1: Validation of the latent correlations within the implicit neural function domain for biomedical data

To further validating the latent correlations within the INR implicit neural function domain for biomedical data, we conducted a statistical analysis to assess the disparities in parameter distributions between samples within the same category and those spanning different categories. To provide a unified evaluation framework, we compared two cross-domain metrics—Structural Similarity Index (SSIM) and the Discrepancy Index within the INR implicit function domain (AWC). This analysis aimed to gauge the disparities between these two metrics with respect to their ability to capture structural and distributional differences in image data. The Discrepancy indices for both metrics are calculated using the following equations:

$$\text{Discrepancy}_{SSIM}=\frac{(SSIM_1-SSIM_2)_{intra-class}}{1-SSIM_{inter-class}}$$

$$\text{Discrepancy}_{AWC}=\frac{(AWC_1-AWC_2)_{intra-class}}{1-AWC_{inter-class}}$$

Here the numerator represents intra-class differences, while the denominator signifies inter-class differences. This implies that when the "Discrepancy" value approaches 0, it signifies reduced discrepancy, indicating a higher degree of domain relevance.

### Supplementary Note 2: Network architecture and network optimization

In our approach, we use a Multi-Layer Perceptron (MLP), consisting of an input layer, several hidden layers and an output layer. For the original data, the corresponding spatial coordinates are first normalized to the [-1,1] range. Then, with the modulation brought by paired cos/sin function, the low-frequency information of coordinates changes across the original data can be converted to high-frequency one in the input layer. The hidden layers comprise a fully connected neural network with Rectified Linear Unit (ReLU) activation functions, designed to emulate the intrinsic structure of data within the high-dimensional embedded space. In contrast, the output layer is a weighted linear combination with bias. Its primary function is to effectively fit the mapping from embedded coordinates to the ultimate intensity.

During network training, the normalized coordinate vectors (2D, 3D, 4D, etc.) are assigned optimization probabilities based on pre-generated saliency maps, where higher probabilities indicate a higher priority in the training optimization process. The entire training process can be regarded as the overfitting of the INR parameter, $f_\theta$, to the input data. For a given input data $x$ and coordinate grid vector $v$, we minimize the objective:

$$arg_{\theta}^{min} \sum_{v \in \mathbb{V}} \mathcal{L}\big(x,\ f_\theta(v)\big)$$

Where $\mathbb{V}$ represents the grid coordinate system that matches the input data, $\mathcal{L}$ measures the difference between original data and INRs output. We use the L2 norm loss as the objective function. In the iterative pursuit of refining network parameters, we employ the Adaptive Moment Estimation Algorithm (Adam)[3], a cutting-edge optimization technique. Additionally, to accommodate the constraints of limited memory resources, we apply the mini-batch gradient descent methodology by sampling a batch of $v$ from $\mathbb{V}$ guided by saliency map.

After completing training on a set of data, due to the high inter-group correlations within pre-grouped data, we employed a weight transfer strategy, preserving the parameters of its hidden layers and using them as the initial parameters for the next set of data. This strategy speeds up the training process and facilitates fast convergence. Following the aforementioned strategy, when we have successfully fitted all the data, we sequentially subtract the network parameters (including weights and biases) corresponding to adjacent group data, encode the residuals using entropy encoding, and ultimately consolidate them into a TAR file, stored in a sequential manner.

More detailed hyper parameters for network training have been listed in the supplementary table 1. We implemented our compression algorithm using PyTorch and CUDA, with all experimental training and testing conducted on NVIDIA GeForce RTX 4090 GPUs.

### Supplementary Note 3: Grouping strategy

For different types of samples, we employed distinct grouping strategies. First, we generated simulated data for three common motion types frequently observed in biomedical data: easing motion, Brownian motion, and periodic motion. Then, each of these datasets was grouped using two distinct strategies: interlace grouping and adjacent grouping. Finally, through a comparative analysis of compression results, we observed that interlace grouping exhibited relatively higher compression performance for simulated data with easing and Brownian motion, whereas adjacent grouping was more effective on simulated data featuring periodic motion. Furthermore, in the context of real biomedical data, we conducted identical experiments for validation and obtained highly similar results. In the case of data containing rapidly moving signals, such as worms exhibiting swift easing motion, we conducted interval sampling along the temporal sequence, generating N groups to ensure the preservation of global motion information within each group. Due to the continuity of frames, there exists a high degree of cross-correlation among these groups. On the contrary, for biomedical sample data exhibiting periodic motion like a beating heart, we grouped the data according to the motion cycles, thereby ensuring the continuous integrity of the cyclic motion.

**Supplementary Table 1. The implementation details of SINCS in the experiments.**

| Experimental data | Network architecture (Number of layers and neurons) | Learning rate | Position encoding frequency |
|---|---|---|---|
| **Large field-of-view 2D images** | 5x64 | 0.001 | $L_x$=10, $L_y$=10 |
| **3D biomedical data** | 5x128 | 0.001 | $L_x$=10, $L_y$=10, $L_t$=6 |
| **Moving *C. elegans* data** | 5x128 | 0.001 | $L_x$=10, $L_y$=10, $L_t$=6 |
| **4D dynamic super-resolution biomedical light-sheet data** | 5x256 | 0.0001 | $L_x$=10, $L_y$=10, $L_z$=6, $L_t$=6 |
| **5D data of multi-channel fluorescently labeled CAR-T cells** | 5x128 | 0.0001 | $L_x$=10, $L_y$=10, $L_z$=6, $L_t$=6 |

**Supplementary Table 2. Comparison of results for grouping strategies for distinct simulated cell data (Easing motion, Brownian motion, and Periodic motion).**

| Motion type | Grouping approach | Network architecture | Compression ratio | Average PSNR | Compression ratio after entropy coding |
|---|---|---|---|---|---|
| Easing motion | Interlace grouping | 4x32 | 1761 | 49.25 | 2001 |
| | Adjacent grouping | 4x32 | 1761 | 47.21 | 1990 |
| Brownian motion | Interlace grouping | 5x96 | 819 | 50.11 | 891.8 |
| | Adjacent grouping | 5x96 | 819 | 49.15 | 891.8 |
| Periodic motion | Interlace grouping | 5x128 | 441 | 46.65 | 480.5 |
| | Adjacent grouping | 5x128 | 441 | 50.38 | 480.9 |

**Supplementary Table 3. Comparison of results for grouping strategies for distinct real dynamic biomedical data (Rapid easing motion and Periodic motion).**

| Data type | Grouping approach | Network architecture | Compression ratio | Average PSNR | Compression ratio after entropy coding |
|---|---|---|---|---|---|
| Worm | Interlace grouping | 5x160 | 582.5 | 47.61 | 642.6 |
| | Adjacent grouping | 5x160 | 582.5 | 46.72 | 631.2 |
| Heart | Interlace grouping | 5x128 | 117 | 41.19 | 133.2 |
| | Adjacent grouping | 5x128 | 117 | 42.74 | 134.4 |

**Supplementary Table 4. The Compression and decompression time comparison of different compression methods.**

| Methods | Compression（seconds） | Decompression（seconds） |
|---|---|---|
| H.264 | 0.832 | 0.443±0.005 |
| H.265 | 2.187 | 0.482±0.005 |
| INR-S | 858.53 | 0.137±0.02 |
| SINCS（Basic） | 518.21 | 0.136±0.02 |
| **SINCS（Weight Transfer）** | 332.49 | |

We took CAR-T cell data as an example and divided it into 30 sub-data sets with each set having a size of 64 x 512 x 512 voxels. The table shows the average compression and decompression time of different compression methods under 260× compression ratio.
